# WDR44 drives de novo α-synuclein aggregation at the lysosomal membrane and promotes neuronal dysfunction in Parkinson’s Disease

**DOI:** 10.64898/2026.04.03.716340

**Authors:** Maxime Teixeira, Razan Sheta, Morgan Bérard, Christine Insinna, Anne-Laure Mahul-Mellier, Constantin V. L. Delmas, Eva Lépinay, Walid Idi, Alexandra Ricard, Esther Del Cid-Pellitero, Christine Trabolsi, Stéphane Gobeil, Marie-Hélène Canron, Erwan Bézard, Alex Rajput, Francesca Cicchetti, Martin Parent, Edward A. Fon, Nagendran Ramalingam, Ulf Dettmer, Frederic Calon, Christopher Westlake, Hilal A. Lashuel, Abid Oueslati

## Abstract

The aggregation of α-synuclein (α-SYN) into Lewy bodies (LBs) is a central event in the pathogenesis of Parkinson’s disease (PD) and related synucleinopathies^1,2^. Despite significant advances in understanding α-SYN self-assembly, the precise sequence of early aggregation steps has not been directly visualized in living neurons. Here, we use an optogenetic-induced protein aggregation system with a high temporal resolution to monitor the onset of α-SYN assembly in neurons. We found that the initiation and accumulation of α-SYN aggregates occur predominantly at the lysosomal membrane, an event driven by the α-SYN N-terminus and modulated by the membrane-associated adaptor protein WD repeat-containing protein 44 (WDR44). Remarkably, we demonstrate that WDR44 knockdown markedly reduced de novo α-SYN aggregation in both neuronal cultures and in vivo, whereas WDR44 overexpression enhances α-SYN aggregation in PD patient-derived iPSC neurons. Consistent with its potential pathogenic involvement, WDR44 aberrantly accumulates in vivo and in the brains of PD patients, where it colocalizes with LB inclusions. Finally, we show that lysosome-associated α-SYN aggregates compromised lysosomal structure and function, leading to neuronal impairment, a phenotype worsened by WDR44 overexpression, linking early aggregation events to downstream toxicity. Together, these findings reveal the earliest dynamic stages of α-SYN oligomerization in living neurons and identify the WDR44-α-SYN interaction as a promising therapeutic target for reducing α-SYN pathology and enabling early intervention in PD.

## Main

The abnormal accumulation of α-synuclein (α-SYN) into Lewy bodies (LBs) and Lewy neurites is a pathological hallmark of Parkinson’s disease (PD) and related synucleinopathies^2,3^. Under physiological conditions, α-SYN exists in a dynamic and tightly regulated equilibrium between a soluble, natively unfolded cytosolic state and a membrane-bound α-helical conformation, primarily associated with synaptic vesicles^4–7^. In PD, however, α-SYN undergoes a conformational shift that promotes its misfolding and self-assembly into insoluble aggregates that progressively evolve into LBs^3,8,9^.

Mounting evidence indicates that the early stages of α-SYN aggregation, particularly the formation of oligomeric species, are major contributors to α-SYN-induced toxicity and the pathogenesis of PD^10,11^. These oligomers interact with lipids, disrupt membrane integrity, and ultimately trigger neuronal dysfunction both *in vitro* and *in vivo*^12–16^. However, the cellular determinants and sequence of events leading to the formation of α-SYN oligomers remains poorly understood, as most insights have emerged from *in vitro* studies relying on oligomers obtain from recombinant α-SYN preparations from cell-free systems^17–19^. Moreover, current experimental models and clinicopathological observations provide only a static, end-stage perspective of α-SYN aggregation, hindering the study of its dynamic oligomerization process.

To overcome these limitations, we previously developed and validated a light-inducible protein aggregation (LIPA) system, which allows precise control of α-SYN aggregation in both cell culture and *in vivo*^20,21^. This approach enables the spatiotemporal induction of α-SYN assemblies and oligomerization leading to the formation of LB-like inclusion that recapitulates many key biochemical hallmarks of authentic LBs^20^, such as phosphorylation at S129, accumulation of p62 and β-sheet structures, and allows the tracking of early interactions with organelles^22^ and protein partners^23^, thereby providing a powerful tool to dissect the initial events of α-SYN aggregation and their pathogenic consequences.

In this study, we combined the LIPA system with advanced imaging techniques and biochemical analysis to monitor and characterize the sequence of events occurring within seconds to minutes of α-SYN structural conversion from monomeric to oligomeric forms in living cells and *in vivo*. Our analysis revealed that aggregation is primarily initiated at the lysosomal membrane, an event essential for the process, as truncations or mutations that disrupt α-SYN-lysosomal interactions drastically alter this process. Moreover, we identified the WD-repeat protein WDR44 as a novel α-SYN interactor that facilitates both the initiation and progression of α-SYN aggregates in two *de novo* α-SYN aggregation models, the LIPA and the 3K-α-SYN systems^24–26^. Notably, WDR44 was also found to accumulate in the brains of PD patients, where it colocalizes with α-SYN within LBs, further supporting its pathological role. Finally, we characterized the functional consequences of α-SYN aggregation at the lysosomal membrane, demonstrating its capacity to drive lysosomal dysfunction and neuronal dysregulation, and showed that WDR44 overexpression amplifies this aggregation-induced impairment. Together, these findings shed light on the earliest molecular and spatial events driving α-SYN aggregation, refining our understanding of the mechanisms that initiate PD pathology and guiding future therapeutic development.

## Results

### Initiation of α-SYN aggregation primarily occurs at the lysosomal membrane in living cells

Leveraging the temporal resolution of the optogenetic LIPA system^20^, we combined live confocal imaging with light-induced α-SYN clustering to monitor, in real time, the formation of α-SYN aggregates in COS-7 cells. Capitalizing on α-SYN well-characterized affinity for lipidic membranes^17,27–29^, we investigated potential interactions between light-induced α-SYN aggregates and various intracellular organelles, including early endosomes (GFP-RAB5B), late endosomes (GFP-RAB7), lysosomes (LAMP1-mGFP), and mitochondria (OTC-GFP). Upon blue light stimulation, we observed a rapid and selective formation of mCherry-positive foci at the LAMP1+ lysosomal membrane (**Fig. 1a**). These nascent aggregates progressively accumulated and increased in size with continued light exposure, suggesting that the lysosomal surface serves as a platform for both the initiation and progressive buildup of α-SYN aggregates. Furthermore, quantification of α-SYN aggregates localized at the lysosomal membrane confirmed this specific and selective interaction, as association with other organelles occurred at significantly lower frequencies (**Fig. 1b; Extended Data Fig. 1a-c**). Notably, kymograph analyses revealed that, once α-SYN aggregates during their trafficking for prolonged periods, indicating a stable interaction (**Fig. 1c, Supplementary Video 1**). Such interaction was not observed with other organelles, further supporting the specificity of the α-SYN-lysosome association **(Extended Data. Fig. 1d-f, Supplementary Videos 2-4**). This α-SYN-lysosome co-trafficking appeared to facilitate the contact and the merging of smaller α-SYN clusters into larger aggregates, suggesting a mechanistic link between lysosomal mobility and the dynamics of α-SYN aggregate formation (**Supplementary Video 1**). The specificity of the α-SYN-lysosome interaction was further confirmed using two controls: LIPA-EMPTY (lacking α-SYN) and LIPA-TDP-43 (amyotrophic lateral sclerosis-associated aggregating protein), neither of which showed significant lysosomal association (**Extended Data. Fig. 1g-i**).

**Figure 1:**
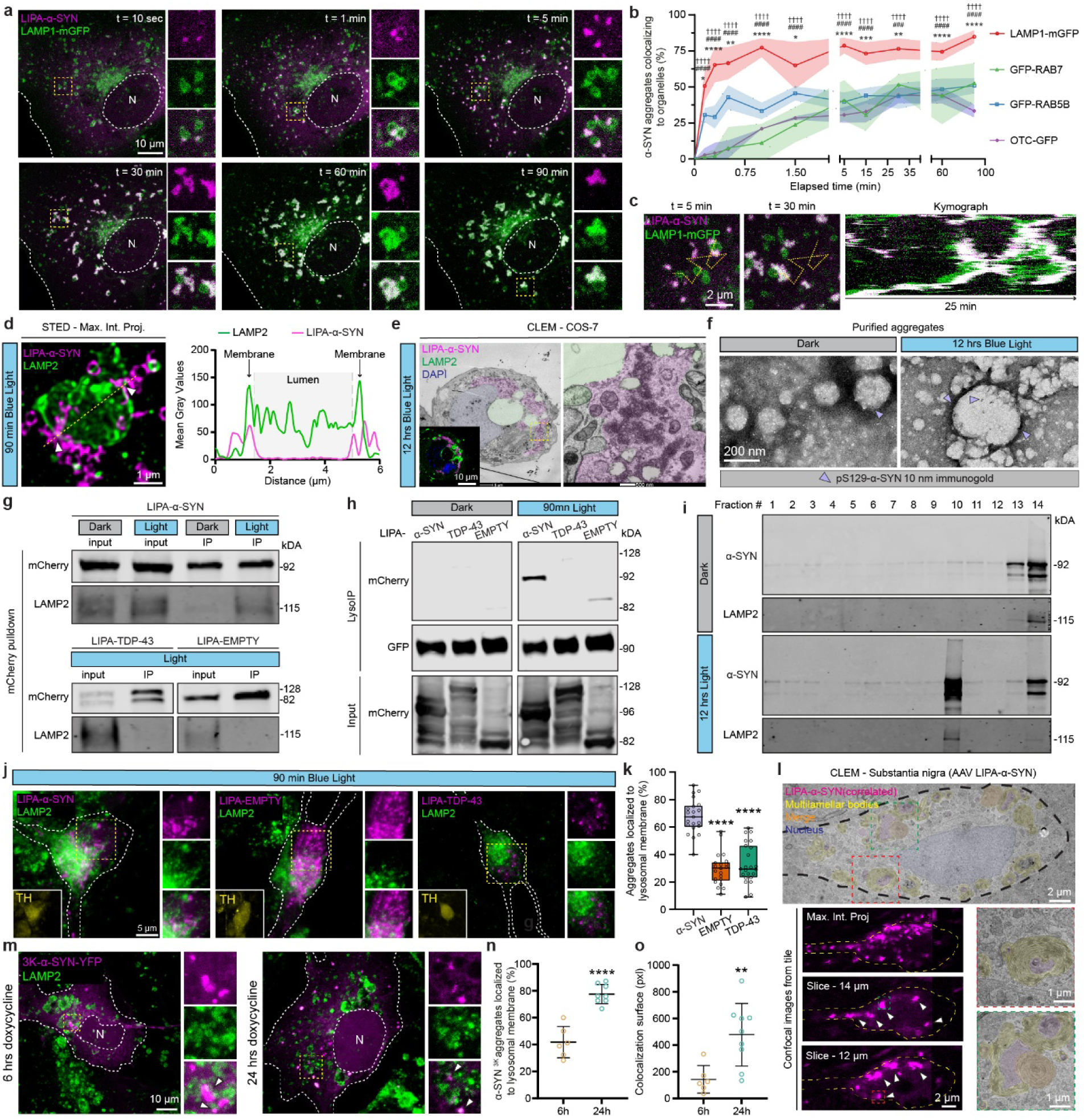
Initiation of **α**-SYN clustering at the lysosomal membrane in different models of Parkinson’s disease. **a**, Representative live-cell confocal microscopy images of COS-7 cells expressing LIPA-α-SYN (magenta) and LAMP1-mGFP (green), showing the rapid α-SYN accumulation at the lysosomal membrane upon blue light exposure (after 10 sec, 1, 5, 30, 60 and 90 min). The small panels on the right show magnified views of the region highlighted by the yellow box in the large panel for each time point. Scale bar, 10 µm. N = nucleus **b**, Quantification of the proportion of LIPA-α-SYN aggregates colocalizing with organelles overtime, displayed as mean ± SEM (n=4, 2-way ANOVA with Dunnett’s multiple comparisons test comparing Lamp1-mGFP with GFP-RAB5B (*), GFP-RAB7 (#) and OTC-GFP (†)). **c**, Representative kymograph over the yellow dot line (left panel) illustrating the co-traffic and the stable interaction between LIPA-α-SYN aggregates (magenta) and lysosomes (green) in COS-7 cells during the first 25 min of blue light exposure (from min 5 to min 30 post-initiation of aggregation). Scale bar, 2 µm. **d**, High-resolution STED microscopy image revealing the localization of LIPA-α-SYN aggregates (magenta) at the lysosomal membrane (green) in COS-7 cells. White arrows are showing focal contact sites. Fluorescence profiles are plotted over the yellow line to show the membrane localization (right panel). Scale bar, 1 µm. **e**, Representative correlative light-electron microscopy (CLEM) image (left panel) and magnified view of the region outlined by the yellow box (right panel) from COS-7 cells overexpressing LIPA-α-SYN (magenta), exposed to 12 hrs of blue light and showing endogenous lysosomal (green) localization at the aggregate’s boundaries. α-SYN aggregates identified by fluorescent microscopy correlated precisely with lysosomal structures in electron microscopy (inset in the lower right corner). Scale bar, 500 nm (TEM), 10 µm (confocal). **f**, Representative TEM micrographs after an immunogold staining (pS129) confirming α-SYN enrichment at the lysosomal membrane (gold particles marked by arrows) from HEK293T purified aggregates. Scale bar, 200 nm. **g**, Representative immunoprecipitation (IP) of mCherry aggregates showing specific biochemical accumulation of lysosomes (LAMP2) with LIPA-α-SYN aggregates. Controls include LIPA-EMPTY and LIPA-TDP43 constructs to confirm IP specificity. **h**, Representative LysoIP assay using streptavidin beads, showing the accumulation of LIPA-α-SYN aggregates in the lysosomal fraction. **i**, Representative Western blots from sucrose gradient fractionation assays, demonstrating LIPA-α-SYN aggregates and the lysosomes co-sedimentation in fraction 10 (F10) upon blue light exposure. **j**, Representative confocal microscopy images of iPSC-derived human dopaminergic neurons (iDA) expressing LIPA-α-SYN, confirming lysosomal interaction. The small panels on the right show magnified views of the region highlighted by the yellow box in the right panel. Inset in the lower right corner shows the TH staining. Scale bar, 5 µm. **k**, Quantitative analysis of LIPA aggregate-lysosome association in iDA neurons, displayed as box and whiskers (n=3, ∼25 cells quantified, one-way ANOVA Tukey’s multiple comparisons test comparing LIPA-α-SYN to the other conditions). **l**, Representative CLEM reconstitution from a mouse dopaminergic neuron overexpressing LIPA-α-SYN (magenta) and exposed to blue light. Correlated LIPA-α-SYN staining colocalizes with multilamellar bodies (manually colored in yellow). Scale bar, 2 µm (Confocal), 2 µm (large TEM), 1 µm (magnified TEM). **m**, Representative confocal microscopy images of COS-7 expressing 3K-α-SYN-YFP (magenta) and stained for endogenous LAMP2 (green). Cells showing accumulation of α-SYN at the lysosomal membrane as early as 6 hrs post-induction of expression, with further increase after 24 hrs. The small panels on the right show magnified views of the region highlighted by the yellow box. Scale bar, 10 µm. **n**, Quantitative analysis of 3K-α-SYN-YFP aggregate-lysosome association presented as mean ± SD, showing an early aggregate-lysosome interaction that increases over time (n=3, ∼10 cells quantified, unpaired t-test). **o**, Quantitative analysis of the lysosomal co-localization surface with the aggregates represented as mean ± SD (n=3, ∼10 cells quantified, unpaired t-test. Each experiment was independently replicated at least three times, and results were reproducible across replicates. *\* p* < 0.05, *\*\* p* < 0.01, *\*\*\* p* < 0.001, *\*\*\*\* p* < 0.0001.

Next, we confirmed α-SYN aggregates localization at the lysosomal membrane using ultrastructural analysis. Super-resolution Stimulated Emission Depletion (STED) microscopy and correlative light-electron microscopy (CLEM) in COS-7 cells revealed that α-SYN aggregates closely associate with the lysosomal surface, without penetrating the lysosomal lumen (**Fig. 1d, e**). Moreover, Transmission Electron Microscopy (TEM) analysis, using anti-phospho-Ser 129 (pS129) specific immunogold labeling of α-SYN aggregates extracted from HEK293T lysates exposed to blue light stimulation for 12 hours (hrs), showed a marked enrichment of gold particle-immunolabeled aggregates localized to the membranes of lysosomal-like vesicles (**Fig. 1f**). Of note, interaction with lysosomes does not appear to promote α-SYN degradation via the autophagy lysosomal pathway (ALP), as total LIPA-α-SYN protein levels remained unchanged upon pharmacological ALP inhibition (**Extended Data. Fig. 1j, k**), suggesting that the lysosomal membrane serves as a site for α-SYN aggregation rather than a facilitator of α-SYN clearance.

To further examine the physical interaction between α-SYN aggregates and lysosomes, we performed co-immunoprecipitation (co-IP) assays. Pull-down using an anti-mCherry antibody revealed a co-IP of the lysosomal marker LAMP2 with α-SYN aggregates, but not with monomeric α-SYN or the control constructs LIPA-EMPTY and LIPA-TDP-43 (**Fig. 1g**). A complementary approach using LysoIP, which allows specific pull-down of intact lysosomes^30,31^, also confirmed a marked recovery of lysosomes with α-SYN aggregates, but not the control constructs (**Fig. 1h**). We additionally confirmed α-SYN-lysosome physical associations under native conditions through sucrose gradient fractionation, a method that assesses protein-protein interactions by analyzing their co-sedimentation profiles in a sucrose gradient^32,33^. In the dark, α-SYN was detected in fractions 13 and 14, while LAMP2 was predominantly present in fraction 14 (**Fig. 1i, top panels**). Upon blue light stimulation, α-SYN and LAMP2 showed a marked co-sedimentation in fraction 10 of the gradient, indicative of the formation of a stable complex between α-SYN aggregates and lysosomes (**Fig. 1i, bottom panels)**.

We then validated α-SYN-lysosome interactions in a neuronal environment, using iPSC-derived human dopaminergic neurons (iDA)^34,35^. Upon blue light stimulation, approximately 70% of α-SYN aggregates co-localized with LAMP2 lysosomal structures inside the soma, whereas LIPA-EMPTY and LIPA-TDP-43 aggregates exhibited minimal lysosomal association, thus confirming α-SYN aggregate-lysosome interactions in human neurons and indicating that this interaction is not cell type-specific (**Fig. 1j, k**). Such interaction was also confirmed *in vivo*, within nigral DA neurons from mice overexpressing LIPA-α-SYN in midbrain and exposed to blue light (1 hr/day) for 7 days (**Extended Data Fig. 1l)**. Interestingly, CLEM analysis revealed α-SYN aggregates associated with concentric membrane layers characteristic of multilamellar bodies (**Fig. 1l**). These structures were previously shown to derive from lysosomes and to retain lysosomal markers in neurons (e.g., LAMP1)^36,37^, confirming that light-induced α-SYN aggregation is also closely associated with lysosome-like compartments *in vivo*. Notably, such multilamellar structures were absent in DA neurons of mice overexpressing LIPA-α-SYN that were not exposed to blue light (**Extended Data Fig. 1m**), supporting the idea that these lysosomal bodies are specifically linked to α-SYN aggregation.

Finally, we showed that the selective interaction of α-SYN aggregates with lysosomes is not unique to the LIPA model and can be generalized to other α-SYN aggregation models. Using COS-7 cells overexpressing α-SYN harboring aggregation-prone mutations E35K, E46K, and E61K (3K-α-SYN), under the control of doxycycline ^24^, we observed a close interaction between lysosomes and newly formed 3K-α-SYN aggregates upon doxycycline induction (**Fig. 1m-o**). These results confirm prior observations of 3K-α-SYN-lysosomes interactions^26,38^, and support the broader notion that the lysosomal membrane might represent a critical site for *de novo* α-SYN aggregation, beyond the LIPA system.

Collectively, our data demonstrate that the structural conversion of monomeric α-SYN into oligomeric aggregates selectively occurs at the lysosomal membrane, identifying this organelle as a primary cellular platform for α-SYN aggregation in living cells.

### Interaction with the lysosomal membrane is essential for the initiation of LIPA-**α**-SYN oligomerization in living cells

Next, we investigated the importance of interaction with the lysosomal membrane in the initiation and progression of α-SYN oligomerization. To this end, we modulated α-SYN membrane-binding capacity by truncating its N-terminal (N-ter) region, which is known to mediate membrane interactions^39,40^. Specifically, we generated LIPA-α-SYN constructs lacking amino acids (AA) 2-4 (LIPA-α-SYN^Δ2-4^), AA 2-10 (LIPA-α-SYN^Δ2–10^), or AA 2-18 (LIPA-α-SYN^Δ2–18^) (**Fig. 2a**). As controls, we deleted C-terminal residues: AA 104-140 (LIPA-α-SYN^Δ104–140^), AA 115-140 (LIPA-α-SYNΔ^115–140^)^41,42^, AA 121-140 (LIPA-α-SYN^Δ121–140^)^41,43,44^, or AA 136-140 (LIPA-α-SYN^Δ136–140^)^41,42^, which do not affect membrane interactions^42,44–46^ (**Fig. 2a**). Using LysoIP in HEK-293T cells, we confirmed that truncations of the membrane-bound N-terminal region, but not the C-terminal region, strongly disrupted α-SYN-lysosomal interactions both at the monomeric and aggregated states (**Fig. 2b**). Strikingly, N-terminal truncations that disrupt membrane binding also completely abolished light-induced α-SYN aggregate formation, whereas C-terminal truncations had no effect on aggregation propensity (**Fig. 2c-e**). Together, these findings demonstrate that interaction with the lysosomal membrane is a critical step in α-SYN aggregation, and that the structural integrity of the N-terminal membrane-binding region is essential for both lysosomal association and subsequent oligomerization.

**Figure 2:**
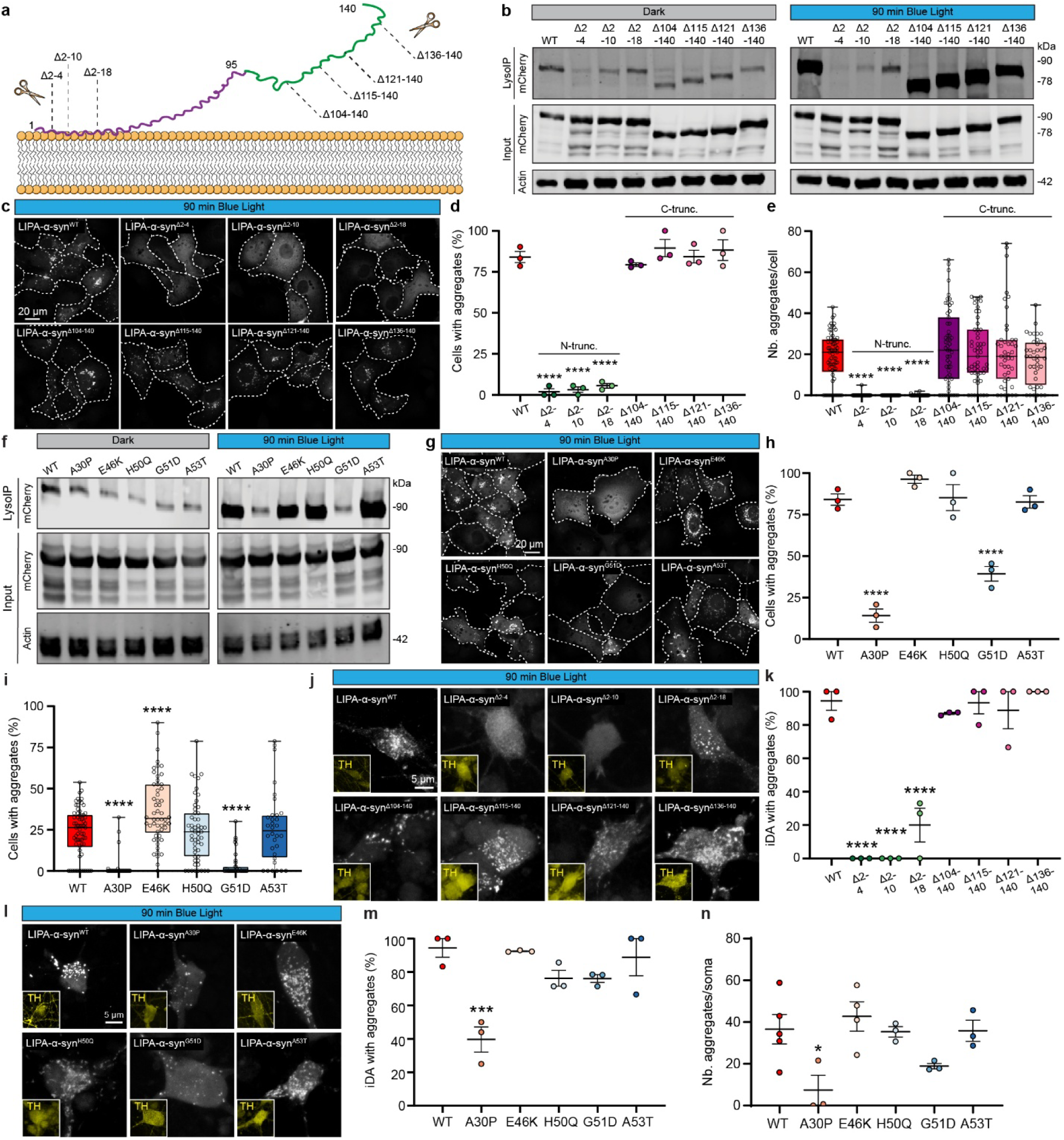
Integrity of **α**-SYN N-terminal region is important for its interaction with lysosomes and aggregation. **a**, Scheme representing the truncations of the LIPA-α-SYN construct: deletions of AA 2-4, 2-10 and 2-18 are directed toward the N-terminal region of α-SYN that binds lysosomal membrane, while deletions of AA 104-140, 115-140, 121-140 and 136-140 are directed towards the free C-tail. **b**, Representative LysoIP Western blot of LIPA-α-SYN truncation variants in the absence (Dark) or presence of light stimulation (90 min). **c**, Representative confocal images of COS-7 cells transfected with truncated LIPA-α-SYN and exposed to the blue light for 90 min. Cell boundaries are marked by white dashed lines. Scale bar, 20 µm. **d, e,** Quantification of the proportion of cells bearing mCherry aggregates (**d,** scatter dot plot ± SEM, n=3, ∼80 cells quantified, one-way ANOVA, Šídák’s multiple comparisons test) and number of aggregates per cells (**e,** box and whiskers, n=3, ∼80 cells quantified; one-way ANOVA, Šídák’s multiple comparisons test) from panel c**. f**, Representative LysoIP Western blot of LIPA-α-SYN harboring PD-linked mutations in the absence (Dark) or presence of light stimulation (90 min). **g**, Representative confocal images of COS-7 cells transfected PD-linked mutated LIPA-α-SYN and exposed to the blue light for 90 min. Cell boundaries marked by white dashed lines. Scale bar, 20 µm. **h**, **i,** Quantification of the proportion of cells bearing mCherry inclusions (**h,** scatter dot plot ± SEM, n=3, ∼80 cells quantified, one-way ANOVA, Šídák’s multiple comparisons test) and number of aggregates per cells (**i,** box and whiskers, n=3, ∼80 cells quantified; one-way ANOVA, Šídák’s multiple comparisons test) from panel g. **j,** Representative confocal images of iDA neurons (TH+), transfected with truncated LIPA-α-SYN constructs and exposed to the blue light for 90 min. Scale bar, 5 µm. **k**, Quantification of the proportion of TH^+^ soma bearing mCherry aggregates. Graphs are represented as scatter dot plot ± SEM (n=3, ∼80 cells quantified, one-way ANOVA, Šídák’s multiple comparisons test). **l**, Representative confocal images of iDA neurons (TH+), transfected with mutated LIPA-α-SYN constructs and exposed to the blue light for 90 min. Scale bar, 5 µm. **m**, Quantification of the proportion of TH^+^ soma bearing mCherry aggregates. Graphs are represented as scatter dot plot ± SEM (n=3, ∼80 cells quantified, one-way ANOVA, Šídák’s multiple comparisons test). **n**, Quantification of the number of aggregates per iDA soma. Graphs are represented as scatter dot plot ± SEM (n=3, ∼80 cells quantified, one-way ANOVA, Šídák’s multiple comparisons test). Each experiment was independently replicated at least three times, and results were reproducible across replicates. * p < 0.05, ** p < 0.01, *** p < 0.001, **** p < 0.0001.

We then assessed the impact of PD-linked mutations, namely A30P, E46K, H50Q, G51D, and A53T, on α-SYN aggregation and interaction with lysosomes using the LIPA system. We observed that A30P and G51D, two mutations known to impair membrane binding^47,48^, but not other mutations, drastically reduced α-SYN association with lysosomes, especially under aggregation conditions, as shown by LysoIP assays (**Fig. 2f**), and also significantly decreased α-SYN aggregate formation in cells (**Fig. 2g-i**). These findings further support the hypothesis that impaired membrane binding disrupts α-SYN oligomerization, likely by preventing its initiation at the lysosomal membrane.

To determine whether this effect also occurs in neurons, we extended our analysis to human iDA neurons expressing either N-or C-terminal truncations or PD-linked mutants (**Fig. 2k-o**). Consistent with our observations in mammalian cells, truncations of the N-terminal membrane-binding region markedly reduced α-SYN inclusion formation in iDA neurons (**Fig. 2j,k**). Similarly, the A30P and G51D mutations diminished α-SYN aggregation in iDA neurons, although the effect of G51D was less pronounced than in mammalian cells (**Fig. 2l-n**).

Altogether, our findings demonstrate that the integrity of the membrane-binding N-terminal region of α-SYN is critical for its interaction with lysosomes and the initiation of oligomerization, which establishes lysosomal membrane binding as a limiting and necessary step in light-induced α-SYN aggregation.

### Vesicle-associated protein WDR44 interacts with **α**-SYN and regulates its aggregation at the lysosomal membrane

Prompted by the rapid and selective interaction of α-SYN with the lysosomal membrane during its oligomerization, we sought to identify the factors that confer specificity to this process. Given that other studies have shown that certain proteins can act as tethers to anchor protein complexes to organelle membranes^49–52^, we focused on protein-protein interactions as potential mediators of α-SYN targeting to lysosomes. Consistent with this idea, our prior work using the LIPA system combined with a proximity-based proteomics approach identified several membrane-associated proteins as early interactors of α-SYN aggregates^23^. Together, these findings further support the importance of interactions with membranes and membrane-associated proteins during the early stages of α-SYN oligomerization, suggesting that these membrane-associated proteins may play a role in tethering α-SYN aggregates to the lysosomal membrane. Among the membrane-associated proteins we previously identified^23^, we focused on those functionally or physically linked to lysosomes, and shown to colocalize with LIPA-α-SYN aggregates in both cell culture and iPSC-derived neurons, namely Insulin-like Growth Factor 2 Receptor (IGF2R), Bridge-Like Lipid Transfer Protein Family Member 3A (BLTP3A), and WD Repeat Domain 44 (WDR44) ^23^. To assess the role of these proteins in the initiation and progression of α-SYN aggregation, we stably knocked down their expression in HEK293T cells using shRNAs (**Extended Data. Fig. 2a-i**). Interestingly, only WDR44 knockdown (KD) led to a significant reduction in α-SYN aggregation, as reflected by the marked decrease in the number of cells with mCherry+ inclusions (**Fig. 3a, b**). This inhibitory effect was further validated using filter retardation assay, which showed a pronounced reduction in detergent-resistant α-SYN aggregates in WDR44^KD^ cells at early (1hr 30) and late time points (24 hrs) following blue light stimulation (**Fig. 3c, d, Extended Data. Fig. 2k**).

**Figure 3:**
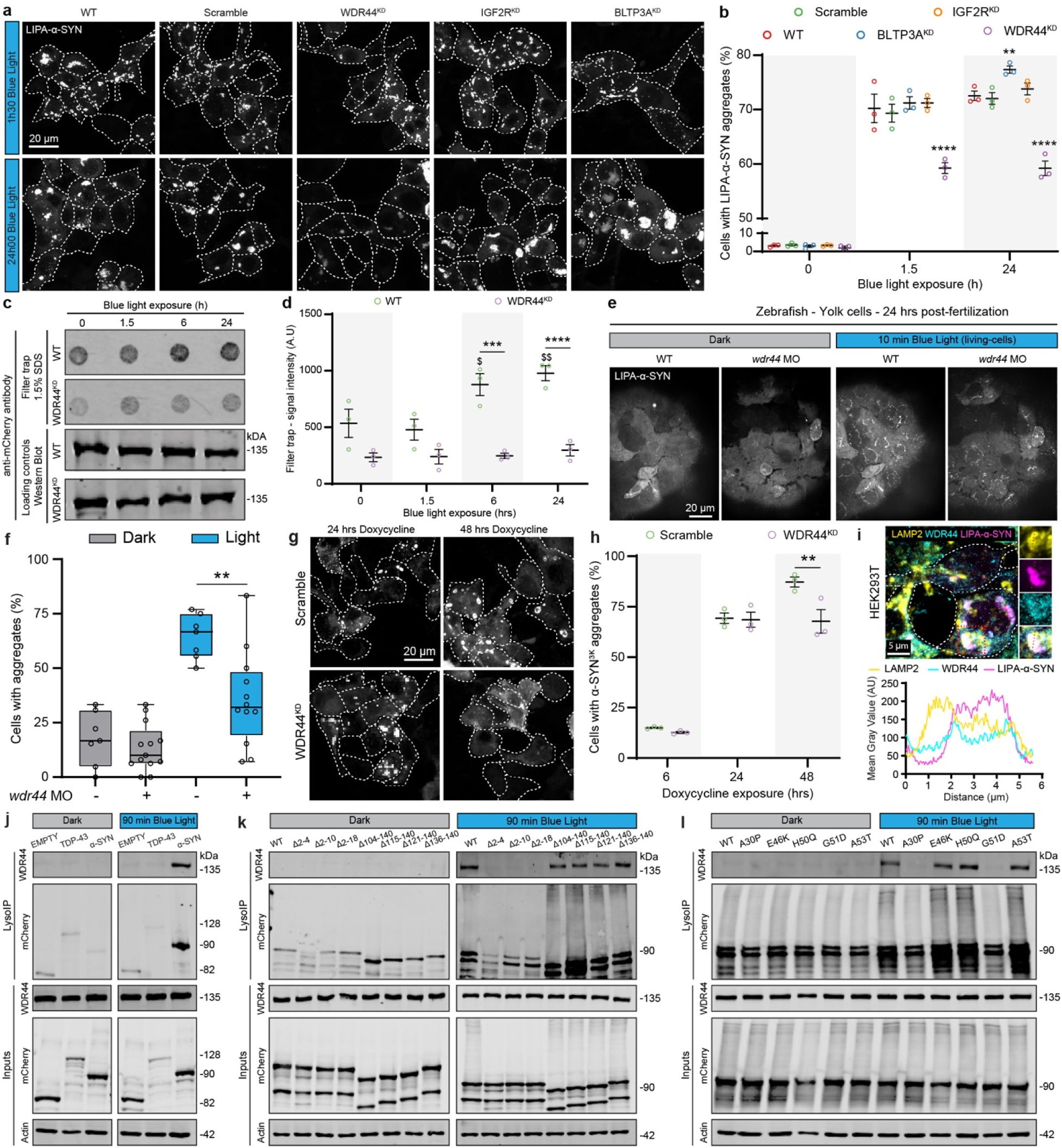
WDR44 interacts with **α**-SYN and modulates its aggregation. **a**, Representative confocal images of HEK293T cells stably expressing shRNAs (scramble, WDR44^KD^, IGF2R^KD^, or BLTP3A^KD^), transiently transfected with LIPA-α-SYN and exposed to blue light for 90 min (top panel) or 24 hrs (bottom panel). White dashed lines delineate individual cells. Scale bar, 20 µm. **b**, Quantification of the proportion of cells bearing LIPA-α-SYN aggregates (scatter dot plot represented as mean ± SEM, n=3, ∼120 cells quantified, one-way ANOVA, Dunnett’s multiple comparisons test). **c**, Filter trap assay of detergent-insoluble α-SYN aggregates from WT and WDR44^KD^ cells transfected with LIPA-α-SYN and exposed to blue light at the indicated timepoints. Representative Western blots for total mCherry levels have been assessed for each condition and shown below as loading controls. **d**, Quantification of filter-trapped aggregates (scatter dot plot represented as mean ± SEM; n=3, two-way ANOVA with Šídák’s multiple comparisons, * compares WT with WDR44^KD^; $ compares each time point to the 0h time point). **e**, Representative live-cell confocal images of yolk sac cells from zebrafish embryos, injected with LIPA-α-SYN mRNA ± *wdr44* morpholino (MO), and stimulated with a green laser. Scale bar, 20 µm. **f**, Quantification of the proportion of cells with aggregates (box and whiskers plot; N=7 to 12 animals per condition; two-tailed unpaired t-test). **g**, Representative confocal images of HEK293T cells stably expressing scramble or WDR44^KD^ shRNAs and transfected with 3K-α-SYN (doxycycline-inducible) and further exposed to 24 hrs or 48 hrs of doxycycline. Scale bar, 20 µm. **h**, Quantification of the proportion of 3K-α-SYN cells bearing aggregates (scatter dot plot, mean ± SEM; n=3, ∼90 cells quantified, two-way ANOVA with Šídák’s multiple comparisons). **i**, Representative confocal image of HEK293T transfected with LIPA-α-SYN (magenta), exposed to 90 min blue light and endogenously stained for WDR44 (cyan) and lysosomes (LAMP2, yellow). The small panels on the right show magnified views of the region highlighted by the red box. The intensity profile from the red-dotted line is displayed below to show the overlap between the three signals. Scale bar, 5 µm. **j**, **k**, **l**, Representative LysoIP membranes comparing the presence or absence of WDR44 in lysosomal fractions in the absence (Dark) or presence of blue light stimulation (90 min). **j**, Western blot membranes of LIPA-EMPTY and LIPA-TDP-43 controls showing specificity of WDR44 pulldown in the presence of LIPA-α-SYN aggregates only. **k**, Representative Western blot membranes of LIPA-α-SYN truncations showing absence of WDR44 in the N-terminal truncated conditions. **l**, Immunoblots of LIPA-α-SYN variants showing absence of WDR44 pulldown in the presence of A30P and G51D mutants. Each experiment was independently replicated at least three times, and results were reproducible across replicates. *\* p* < 0.05, *\*\* p* < 0.01, *\*\*\* p* < 0.001, *\*\*\*\* p* < 0.0001.

To confirm these findings *in vivo*, we used a Wdr44^KD^ zebrafish model overexpressing LIPA-α-SYN. Embryos were injected with *wdr44* morpholinos (MOs) to reduce endogenous Wdr44 expression, validated via a GFP reporter RNA containing the 5’ UTR MO target site as previously described^53^. LIPA-α-SYN mRNA were injected at the one cell stage, and embryos were exposed to blue light 24 hrs post-fertilization to induce aggregation. We observed robust mCherry+ inclusion formation upon light stimulation in control embryos, with more than 70% of yolk sac cells per embryo displaying LIPA-α-SYN aggregates. Interestingly, in *wdr44* MO-injected embryos, the number of cells with inclusions was significantly reduced to ∼30% (**Fig. 3e, f**), supporting the hypothesis of a critical role of Wdr44 in modulating α-SYN aggregation *in vivo*.

To investigate whether the function of WDR44 in modulating α-SYN aggregation extends beyond the LIPA system, we analyzed 3K-α-SYN aggregation in WDR44^KD^-HEK293T cells. Forty-eight hours following doxycycline induction, WDR44 depletion significantly reduced the proportion of cells exhibiting α-SYN inclusions, demonstrating that WDR44 acts as a critical regulator of α-SYN aggregation across distinct experimental models (**Fig. 3g, h**).

In light of a direct interaction with α-SYN at the lysosomal membrane, we therefore performed immunocytochemistry, confirming the colocalization of WDR44 with α-SYN aggregates and lysosomes in HEK293T cells (**Fig. 3i**), which was absent in the conditions transfected with the controls LIPA-EMPTY or LIPA-TDP-43 (**Extended Data Fig. 2l**). This interaction was not restricted to the LIPA system, as we also observed endogenous WDR44 accumulation at the lysosome-aggregate interface when using the 3K-α-SYN model in HEK293T cells (**Extended Data Fig. 2m**). We further confirmed that α-SYN aggregates and lysosomes were also colocalizing with endogenous WDR44 in iDA neurons (**Extended Data Fig. 2n**). *In vivo*, WDR44 was also found to colocalize with α-SYN inclusions in midbrain dopaminergic neurons from LIPA-α-SYN–overexpressing mice exposed to blue light. However, due to suboptimal lysosomal staining in these neurons, we were unable to reliably assess the triple colocalization of WDR44, α-SYN aggregates, and lysosomes in this model (**Extended Data Fig. 2o**).

Next, we confirmed the physical interaction between WDR44, α-SYN, and lysosomes using LysoIP. Notably, we observed a strong recovery of lysosomes with endogenous WDR44 exclusively under aggregation conditions, but not with monomeric α-SYN or controls, indicating a specific and selective recruitment of WDR44 during the α-SYN aggregation process (**Fig. 3j**). This observation was further supported by LysoIP experiments using a truncated (**Fig. 3k**) and PD-linked α-SYN forms (**Fig. 3l**), where endogenous WDR44 pulldown with lysosomes and α-SYN was impaired under conditions that blocked α-SYN-lysosome interaction and prevented LIPA-α-SYN aggregation (N-terminal truncation, A30P and G51D mutations) (**Fig 3k-l**).

Collectively, our results reveal for the first time that WDR44 is an important modulator of α-SYN aggregation across multiple experimental models, both in culture and *in vivo*. Furthermore, WDR44 interacts selectively with α-SYN and lysosomes under aggregation conditions, suggesting that this protein may act as a tether anchoring α-SYN to the lysosomal membrane, a key factor in the aggregation process. This tethering function of WDR44 has also been reported in connecting RAB11 with endolysosomal vesicles and coordinating vesicle trafficking within cells^49,54–56^.

### WDR44 aberrantly accumulates in human PD brains and mouse models, and colocalizes with **α**-SYN within LBs

The colocalization of WDR44 with α-SYN inclusions in culture and *in vivo* and the effect of WDR44 knockdown on α-SYN aggregation across different experimental models raised the question of the pathological relevance of WDR44 in PD pathogenesis. To address this question, we first examined whether WDR44 protein levels are altered in PD. We analyzed *substantia nigra* extracts from PD patients (n=24, age = ∼80) and age-matched controls (n=19, age = ∼75) from the Saskatchewan movement disorders program^57^ (**Table I**). Western blot analysis and quantification revealed significantly higher level of WDR44 protein levels in the PD midbrain compared to controls (**Fig. 4a, b**). Notably, this increase in WDR44 levels positively correlated with the accumulation of pathological, insoluble pS129-α-SYN in PD brains, suggesting a potential link between α-SYN pathology and WDR44 upregulation (**Fig. 4c**). This observation aligns with a recent clinicopathological study reporting a correlation between aberrant accumulation of WDR44 and the presence of pathological α-SYN in the brains of individuals with PD and Parkinson’s disease dementia (PDD)^58^. Consistent with this observation in human brains, increased levels of WDR44 were also observed in dopaminergic neurons of the *substantia nigra* in the Thy1-α-SYN transgenic mouse model of PD, further supporting an association between pathological α-SYN and WDR44 upregulation (**Fig. 4d, e**).

**Figure 4:**
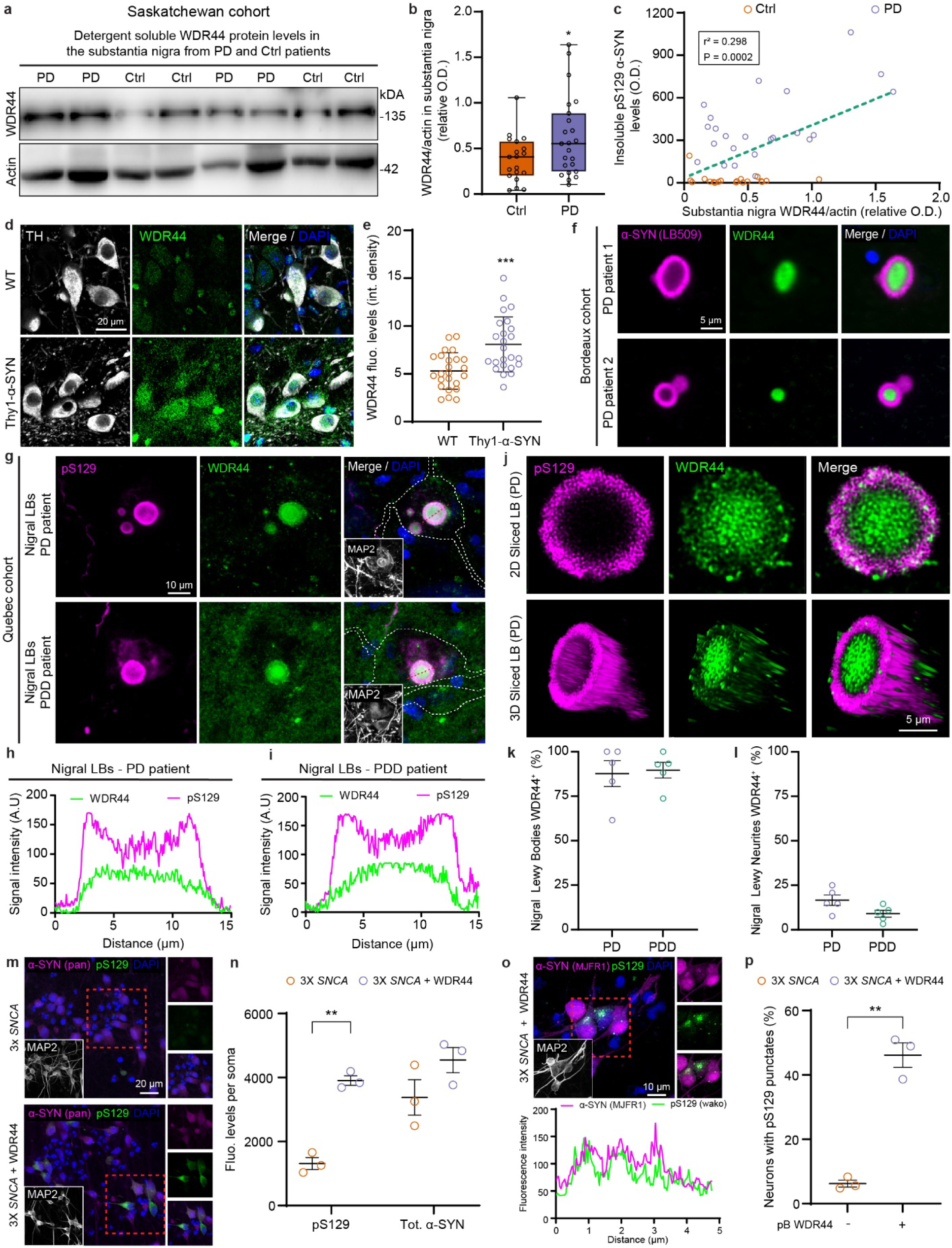
WDR44 protein levels are elevated in PD and PDD patients and accumulate within the core of Lewy Bodies. **a**, Representative Western blot of detergent-soluble WDR44 and actin in substantia nigra (SN) lysates from control (Ctrl) and PD patients (PD) from the Saskatchewan movement disorders program. **b**, Quantification of WDR44 normalized to actin (box and whiskers plot, N=19 Ctrl, N=23 PD; Unpaired t test with Welch’s correction). Three technical outliers were excluded based on ROUT analysis (one from the PD group and two from the control group). **c**, Correlation between WDR44 and pS129 α-SYN levels across individuals (Pearson r² = 0.298, p = 0.0002). **d**, Representative confocal images showing TH+ nigral neurons (gray) stained for WDR44 (green) in WT and 8-month-old Thy1-α-SYN mice. Scale bar, 20 µm. **e**, Quantification of WDR44 fluorescence intensity (protein levels) per neuron showing an increased WDR44 levels in aged Thy1-α-SYN mice (scatter dot plot, mean ± SD; N=5 mice per group; 20-30 cells per group, unpaired t-test). **f**, Representative high magnification epifluorescence images of nigral LBs from two individual PD patients from the Bordeaux cohort stained for human α-SYN (LB509, magenta) and WDR44 (green) showing aberrant accumulation of WDR44 in the core of the LBs. Scale bar, 5 µm. **g**, Representative confocal images of nigral LBs from PD and PDD brains stained for pS129 α-SYN (WAKO, magenta), WDR44 (green), and MAP2 (white). Scale bar, 10 µm. **h**, **i** Corresponding intensity profiles from the PD (h) or PDD patient (i) displayed from the red-dotted line showing WDR44 accumulation in the core of the inclusions. **j**, 2D sliced (top) and 3D reconstructed (bottom) LBs in a PD brain stained for WDR44 (green) and α-SYN pS129 (WAKO, magenta). Scale bar, 5 µm. **k, l** Quantification of WDR44-positive inclusions (in LBs (**k**) or LNs (**l**)) from PD and PDD patients in the *substantia nigra*, showing high enrichment in WDR44-postive LBs for PD and PPD patients (scatter dot plot, mean ± SEM; N=5 patients per condition with ∼20 LBs and ∼70 LNs imaged per patient). **m**, Representative confocal images showing neurons derived from 3X *SNCA* hiPSC cells overexpressing (bottom panel) or not (top panel) WDR44, 21 days *in vitro* post differentiation. The small panels on the right show magnified views of the region highlighted by the red box. Cells overexpressing WDR44 shows increase pS129 fluorescence levels and inclusions formation. Scale bar, 20 µm **n**, Quantification of α-SYN pS129 (phosphorylated protein levels; WAKO) and total α-SYN (pan-syn) total protein levels) per soma (scatter dot plot, mean ± SEM; n=3, ∼75 cells quantified per group; 2-way ANOVA with Šídák’s multiple comparisons test). **o**, Representative confocal images showing neurons derived from 3X *SNCA* hiPSC cells overexpressing WDR44 and further stained with C-terminal α-SYN antibody (MJFR1) and pS129 (WAKO) showing overlap between the two signals within inclusions. The small panels on the right show magnified views of the region highlighted by the red box. Fluorescence intensity profiles (bottom panel) showing the overlap are displayed over the red dotted line in the merge image. Scale bar, 10 µm. **p**, Quantification of cells bearing pS129^+^ inclusions, relative to panel **o** (scatter dot plot, mean ± SEM; n=3, ∼90 cells quantified per replicate and conditions, unpaired t-test with Welch’s correction). All *in vitro* experiments involving 3X *SNCA* hiPSC cells were independently replicated three times. *\* p* < 0.05, *\*\* p* < 0.01, *\*\*\* p* < 0.001, *\*\*\*\* p* < 0.0001.

**Table I.**
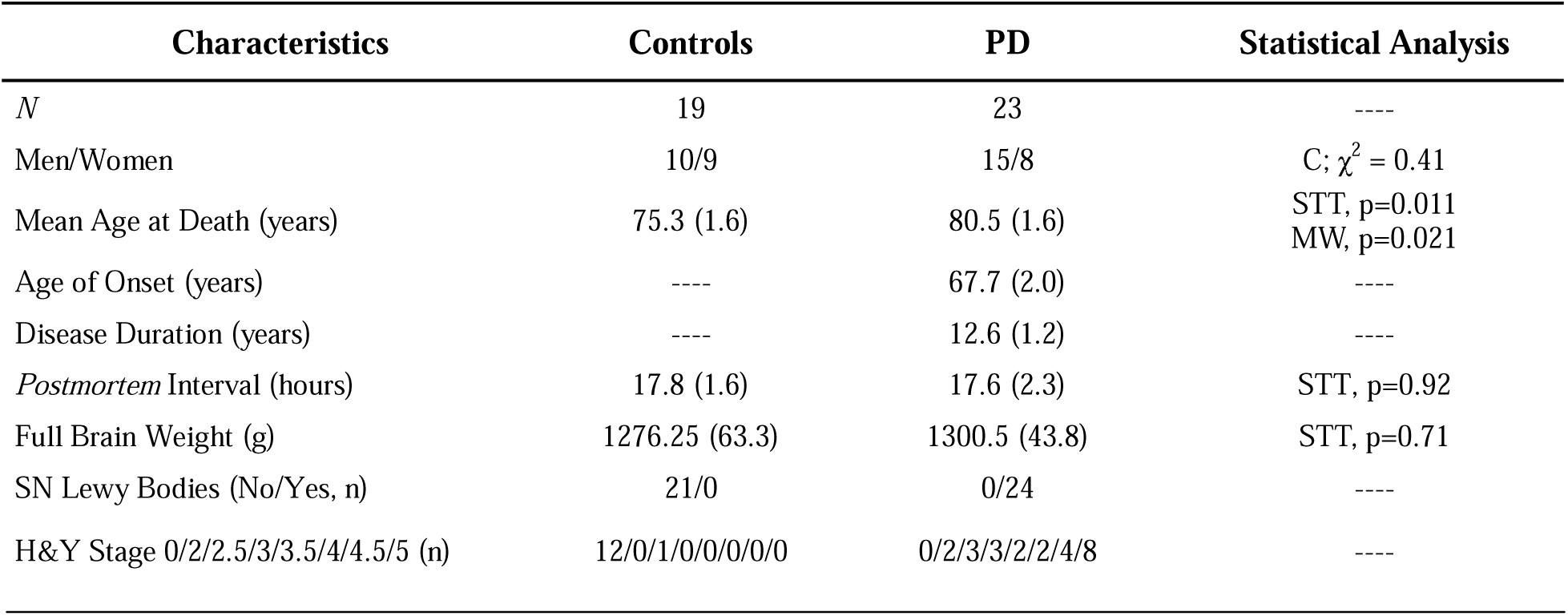
Description of the Saskatchewan cohort used for biochemical assays. The postmortem diagnosis was performed by a neuropathologist for each patient and the clinical assessments were accomplished by AR. The values are presented as means (SD). Statistical analyses were performed using Contingency (C), Student unpaired t-test (STT) or Mann-Whitney tests (MW). Abbreviations: PD, Parkinson’s disease patients; C, contingency; H&Y, Hoehn and Yahr scale; MW, Mann-Whitney; ROD, relative optical density; SD, standard deviations; SN, substantia nigra; STT, Student unpaired t-test.

| Characteristics | Controls | PD | Statistical Analysis |
| --- | --- | --- | --- |
| *N* | 19 | 23 | ---- |
| Men/Women | 10/9 | 15/8 | C; $\chi^2 = 0.41$ |
| Mean Age at Death (years) | 75.3 (1.6) | 80.5 (1.6) | STT, p=0.011<br>MW, p=0.021 |
| Age of Onset (years) | ---- | 67.7 (2.0) | ---- |
| Disease Duration (years) | ---- | 12.6 (1.2) | ---- |
| *Postmortem* Interval (hours) | 17.8 (1.6) | 17.6 (2.3) | STT, p=0.92 |
| Full Brain Weight (g) | 1276.25 (63.3) | 1300.5 (43.8) | STT, p=0.71 |
| SN Lewy Bodies (No/Yes, n) | 21/0 | 0/24 | ---- |
| H&Y Stage 0/2/2.5/3/3.5/4/4.5/5 (n) | 12/0/1/0/0/0/0/0 | 0/2/3/3/2/2/4/8 | ---- |

We next investigated whether WDR44 is present within LBs, the hallmark of PD and related disorders. Double immunofluorescence labeling revealed colocalization of WDR44 with pS129-α-SYN within LBs structure in two independent patient cohorts: the Bordeaux cohort (PD patients, n=2; Table II; **Fig. 4f**) and the Quebec City cohort (PD, n=5; PDD, n=5; Table II; **Fig. 4g**). Further analysis of WDR44 and α-SYN distribution within LBs using intensity profiling (**Fig. 4h, i**) and STED 3D imaging revealed a characteristic ring-like architecture in nigral LBs, with WDR44 immunoreactivity enriched within the core, surrounded by pS129-α-SYN staining (**Fig. 4j**). This immunoreactivity was present in more than 85% of the LB of the SN, regardless of the disease (PD or PDD) (**Fig. 4k**). Interestingly, Lewy neurites were less prone to be colocalized with WDR44, with only around 20% of the inclusions being positive for WDR44 staining, suggesting a mechanism of accumulation that mostly occurs in the cytosol of neuronal somas (**Fig. 4l**).

To investigate the intricate relationship between α-SYN aggregates and WDR44, we mimicked WDR44 overexpression, using piggybac transposons system, in iPSC-derived human neurons (iNeurons) bearing the triplication of *SNCA* gene (3X *SNCA*) (**Extended Data Fig. 3a-b**). After 21 days *in vitro,* we observed that iNeurons overexpressing WDR44 displayed a significant increase of pS129-α-SYN levels, while total α-SYN level remains unchanged (**Fig. 4m, n**). Moreover, we observed the formation of pS129-positive inclusions in 50% of the neurons (**Fig. 4o, p**), suggesting that WDR44 overexpression is sufficient to induce the formation of *de novo* aggregates in PD-derived (3X *SNCA)* neurons.

Collectively, our results suggest a functional link between WDR44 accumulation in PD brains and α-SYN aggregation and point to WDR44 as an important modulator of α-SYN pathology and PD pathogenesis.

**Tab le II.**
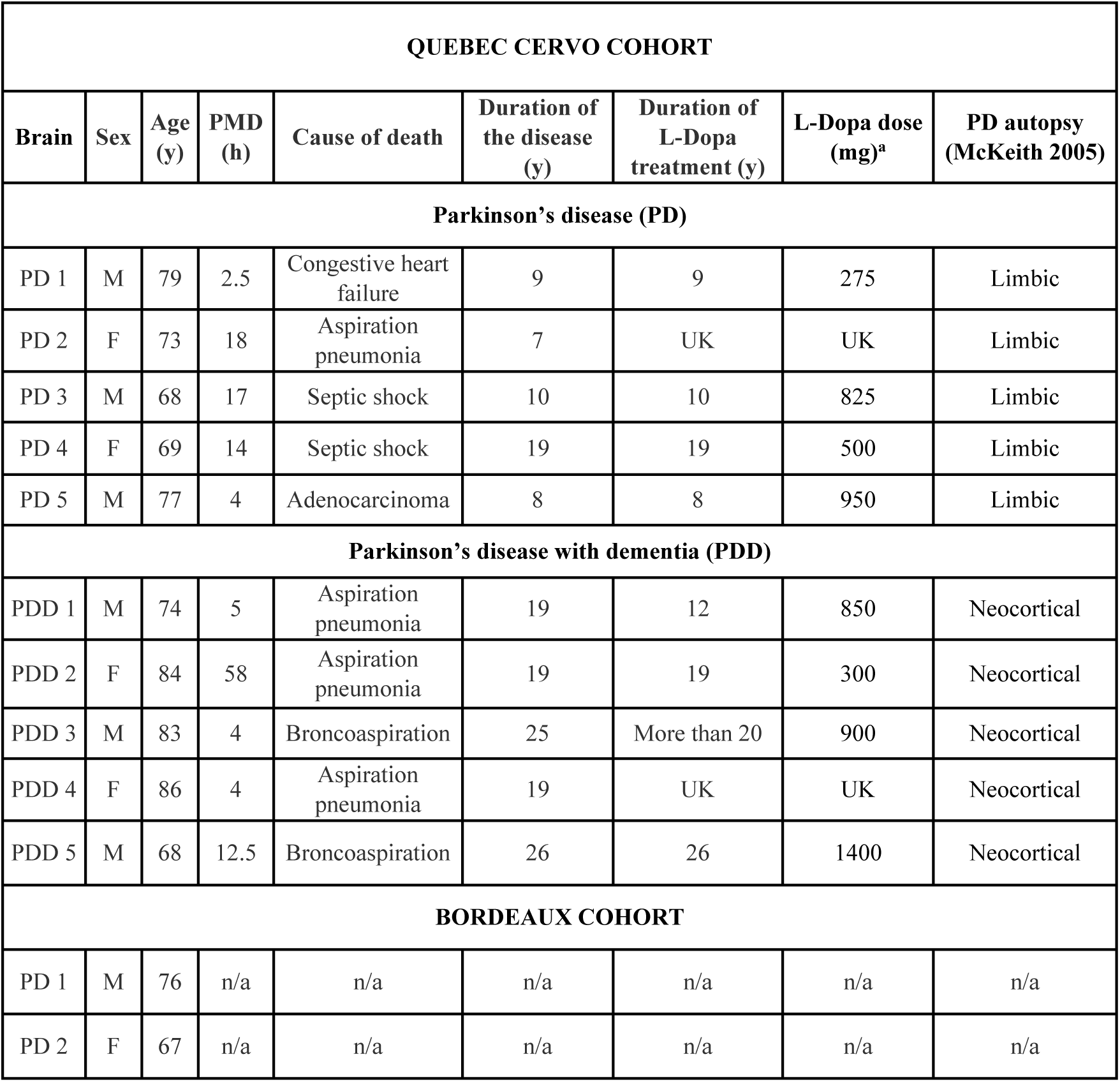
Description of the Que bec cohort and Bordeaux cohort used for immunofluorescence assays.

### WDR44 overexpression exacerbates lysosomal dysfunction and neuronal loss induced by **α**-SYN aggregation at the lysosomal membrane

We subsequently examined the functional consequences of α-SYN aggregation at the lysosomal membrane and assessed how WDR44 overexpression influences this process, aiming to replicate the pathological context observed in PD. Live-cell imaging revealed that the association of α-SYN with lysosomes significantly impaired the mobility of these organelles in the cytoplasm. Tracking individual lysosomes movement during the initial 15 minutes of light stimulation showed numerous lysosomes traveling relatively long distances in multiple directions, as reflected by a heatmap displaying the distance traveled by each lysosome. However, with prolonged light exposure, lysosomal movement was notably reduced, with fewer organelles in motion and shorter travel distances, thus reflecting lysosomal motility deficits (**Fig. 5a**). Quantitative analysis confirmed a significant decrease in both the total distance traveled and the number of motile lysosomes (**Fig. 5b-c**). These findings indicate that α-SYN aggregation at the lysosomal membrane disrupts lysosomal trafficking, a hallmark of lysosomal dysfunction.

**Figure 5:**
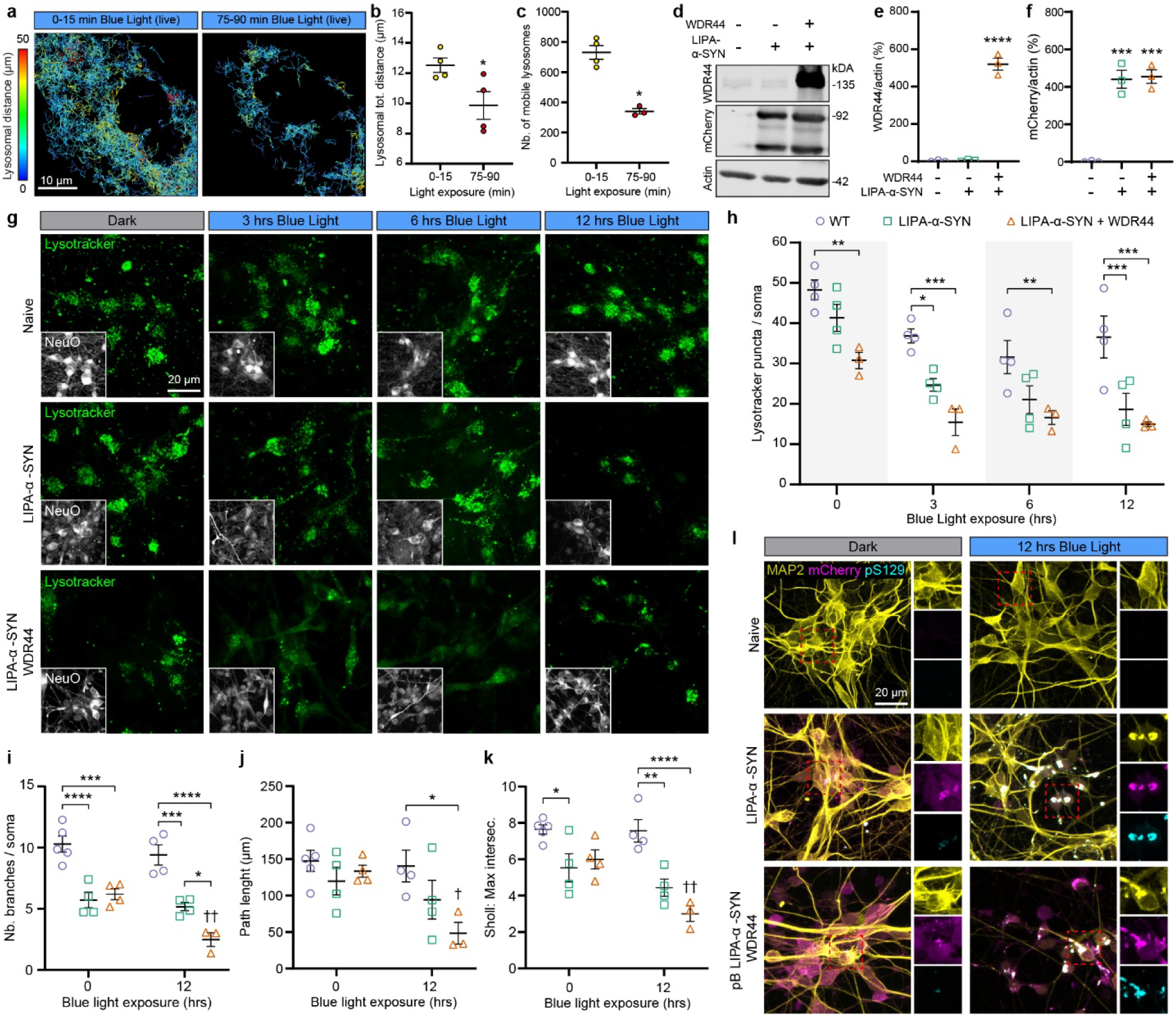
WDR44 overexpression exacerbates **α**-SYN-induced lysosomal dysfunction and neuronal impairment. **a**, Representative lysosomal traces from COS-7 cells transfected with LIPA-α-SYN and Lamp1-mGFP exposed to blue light; recorded in real time for 1h30. Weka segmentation was performed, and individual traces were generated using Trackmate 7 to visualize individual lysosomal movements. Tracked lysosomes are color-coded to indicate the distance traveled within the first and the last 15 min of the recording (red, long distance; blue, short distance). Scale bar = 10 µm. **b**, Quantification of the total distance traveled by individual lysosomes in living cells in the first (0-15 min) and the last (75-90min) 15 min of acquisition (Scatter dot plot, mean ± SEM; n=4, ∼800 lysosomes tracked per cell; paired t-test), **c**, Quantification of the number of individual lysosomes tracked in the first (0-15 min) and the last (75-90 min) 15 min of acquisition (Scatter dot plot, mean ± SEM; n=4, ∼800 lysosomes tracked per cell; paired t-test), **d**, Representative Western blot confirming piggyBac-driven LIPA-α-SYN and WDR44 overexpression in iPSC-derived neuronal cultures (with endogenous WDR44 for reference). **e**, Quantification of WDR44 protein levels normalized to actin (scatter dot plot, mean ± SEM; n=3; One-way ANOVA with Tukey’s multiple comparisons test). **f**, Quantification of LIPA-α-SYN protein levels normalized to actin (scatter dot plot, mean ± SEM; N=3; One-way ANOVA with Tukey’s multiple comparisons test). **g**, Representative confocal images of DIV14 neurons stably expressing no construct (naive), LIPA-α-SYN, or LIPA-α-SYN + WDR44. Differentiated neurons were exposed to blue light at different time points (3, 6 and 12 hrs) and imaged in real-time using LysoTracker (green) and NeuO (white) staining. A decrease of LysoTracker punctate was observed in LIPA-α-SYN and LIPA-α-SYN + WDR44 overexpression conditions. Scale bar, 20 µm. **h**, Quantification of LysoTracker fluorescent punctate per neuronal soma, showing a decrease in LIPA-α-SYN condition aggravated with WDR44 overexpression (Scatter dot plot, mean ± SEM; n=4, ∼ 50 cells per replicate (5 to 7 fields of view); 2-way ANOVA with Tukey’s multiple comparisons test). **i**, Quantification of the total number of branches per neuron analyzed using the SNT plugin from ImageJ using MAP2 signal. Significant decrease is observed in LIPA-α-SYN condition, aggravated with WDR44 overexpression (Scatter dot plot, mean ± SEM; n=4, ∼ 20 cells per replicate; 2-way ANOVA with Šídák’s multiple comparisons test). **j**, Quantification of the neurite length per neuron analyzed using SNT plugin from ImageJ using MAP2 signal. Significant decrease is observed in LIPA-α-SYN + WDR44 overexpression (Scatter dot plot, mean ± SEM; n=4, ∼ 20 cells per replicate; 2-way ANOVA with Šídák’s multiple comparisons test). **k**, Quantitative sholl analysis of the maximum intersection number of the arborescence using SNT plugin from ImageJ using MAP2 signal. Significant decrease is observed in LIPA-α-SYN condition, aggravated with WDR44 overexpression (Scatter dot plot, mean ± SEM; n=4, ∼ 20 cells per replicate; 2-way ANOVA with Šídák’s multiple comparisons test). **l**, Representative confocal images from the cells analyzed in **i**, **j**, **k**. Neurons were immunostained for MAP2 (yellow) and pS129-α-SYN (cyan, Abcam). The small panels on the right show magnified views of the region highlighted by the red box.. Cells bearing pS129^+^ inclusions show a disrupted neuronal arborescence. Scale bar, 20 µm. Each experiment was independently replicated at least three times, and results were reproducible across replicates. *\* p* < 0.05, *\*\* p* < 0.01, *\*\*\* p* < 0.001, *\*\*\*\* p* < 0.0001.

**Figure 6:**
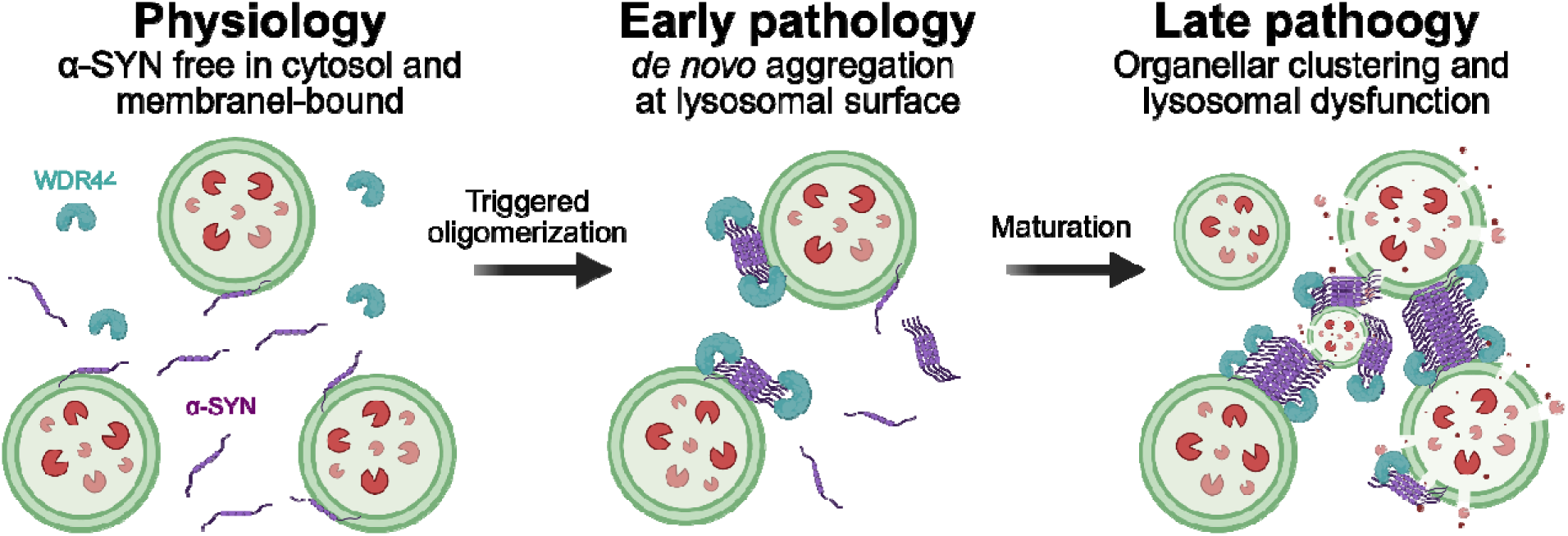
WDR44 interacts with **α**-SYN aggregates at the lysosomal membrane, promoting organelle clustering and impairing lysosomal function. Schematic model of the mechanistic sequence of events surrounding WDR44-α-SYN interplay at the lysosomal membrane. Left panel: physiologically, α-SYN (purple) and WDR44 (cyan) remain in the cytosol, while part of monomeric α-SYN is associated with the lysosomal membrane (green vesicles). Middle panel: Once aggregation is triggered, WDR44 binds to nascent α-SYN aggregates and promotes membrane-proximal nucleation and elongation of the aggregates, stabilizing their association with lysosomes. Right panel: Over time, growing aggregates bridge neighboring lysosomes and accumulate WDR44 in the vicinity of the inclusions, leading to a feed-forward loop inducing organelle clustering and lysosomal membrane permeabilization connected to cell death. Scheme not to scale.

We then assessed the functional impact of α-SYN aggregation on lysosomal activity, in the context of WDR44 overexpression in a disease-relevant model to mimic the environment observed in PD and PDD patients. Thus, we transduced our iPSC line bearing the Ngn2 with piggyBac transposons that allow for stable LIPA-α-SYN and/or WDR44 overexpression under a tetracycline-inducible promoter activated only after neuronal differentiation (**Fig. 5d-f)**. PiggyBac transposons induced a strong and comparable expression of LIPA-α-SYN in both cell lines and WDR44 was highly overexpressed in the line mimicking the PD overexpression phenotype up to 500-fold as compared to the endogenous protein levels (**Fig. 5e, f**). Combining those differentiated neurons with LysoTracker, a dye that accumulates specifically in acidic compartments and serves as a proxy for active lysosomes, we observed a significant decrease in its signal intensity with prolonged light-induced α-SYN aggregation inside living neurons, reflecting impaired lysosomal acidification and function as opposed to the control cells (**Fig. 5g, h**). Notably, overexpression of WDR44, mimicking its pathological accumulation observed in PD, further exacerbated lysosomal dysfunction, leading to an even more pronounced reduction in LysoTracker signal (**Fig. 5g, h**).

Finally, we evaluated the effect of α-SYN aggregation on neuronal health. Neurons exposed to prolonged light stimulation and α-SYN aggregation exhibited clear signs of morphological impairment, including a reduction in the number of branches, neurite length and the neuronal arborescence complexity (**Fig. 5 i-l**). Importantly, WDR44 overexpression further intensified these early signs of neurodegeneration, suggesting that, as observed in PD brains, WDR44 contributes to the worsening of α-SYN-induced neuropathology (**Fig. 5i-l**).

Together, these findings indicate that α-SYN aggregation at the lysosomal membrane impairs lysosomal function and trafficking, and that WDR44 overexpression exacerbates these defects, ultimately leading to enhanced neuronal degeneration, mirroring key aspects of PD pathology.

## Discussion

The early dynamics of α-SYN aggregation in living cells remain poorly understood, largely due to the lack of suitable tools and model systems. In this study, we combined our newly developed optogenetic α-SYN aggregation model with advanced live-cell imaging to visualize α-SYN *de novo* aggregation directly. We found that this process primarily occurs at the lysosomal membrane and depends on the integrity of the N-terminal membrane-binding region of α-SYN, as well as its interaction with WDR44, which anchors nascent oligomers to lysosomal surfaces. Post-mortem analyses unveiled that WDR44 aberrantly accumulates in PD brains and colocalizes with α-SYN within LBs, establishing a clear relevance of this protein to PD pathogenesis. Functionally, the occurrence of α-SYN aggregation at the lysosomal membrane impairs the integrity and the function of this organelle, leading to neuronal dysfunction; effects that are further exacerbated by WDR44 overexpression. These findings provide new insights into the early dynamic processes in the formation of α-SYN aggregates and identify WDR44 as a key mediator of α-SYN aggregation and LB formation.

A key finding of this study is the early and specific association of α-SYN with the lysosomal membrane during its structural conversion from monomeric to aggregated forms. Captured here with second-scale temporal resolution in living cells, this event appears to be unique to α-SYN, is reproducible *in vivo*, and is also observed in an independent *de novo* α-SYN aggregation model, 3K-α-SYN cells^26,38^, suggesting that engagement with the lysosomal membrane represents a general and early step in *de novo* α-SYN aggregation. This interaction is consistent with the growing body of literature linking α-SYN and lysosomes. Ultrastructural analyses of LBs in human brains, as well as LB-like inclusions in experimental models, reveal that they are largely composed of vesicular structures resembling lysosomes and autophagosomes^11,20,59^. Moreover, post-mortem studies have identified lysosome-associated LBs in PD brains^44,60^, reinforcing the pathological relevance of α-SYN-lysosome interactions.

The temporal and real-time analyses further reveal that the lysosomal interaction is critical not only for the initiation of aggregation but also for aggregate buildup and growth. Live-cell imaging and kymograph analyses show rapid α-SYN accumulation on the lysosomal membrane within seconds, followed by prolonged and stable association. By hijacking lysosomal transport and fusion/fission dynamics, α-SYN exploits these organelle processes to connect small cytosolic aggregates, thereby facilitating the formation of larger assemblies. Such a strategy is reminiscent of other proteins and biological entities that utilize organelle surfaces to achieve physiological or pathological functions, like RNA granules that hitchhike on lysosomes for long-distance transport^61^ or peroxisomes^62,63^ and lipid droplets^63^ that hitchhike on early endosomes.

The close engagement of nascent α-SYN aggregates with lysosomes underscores the pivotal role of α-SYN/lysosomal membrane interactions in driving aggregation and suggests that interaction is an important event; if disrupted, this interface may attenuate aggregate formation. Substantial evidence indicates that α-SYN associates with membranes through its N-terminal region, which can adopt either a continuous α-helical conformation (AA 1-95) or a broken double helix separated by a short linker around Glycine 41^28,64^. These structural states are strongly influenced by the lipid composition of the target vesicles ^65–67^. Accordingly, perturbing the integrity of this membrane-binding domain would be expected to weaken lysosomal association and suppress aggregation. Our findings support this prediction: N-terminal truncations (Δ2-4, Δ2-10, Δ2-18) markedly reduced lysosomal interaction, as shown by LysoIP, and completely abolished α-SYN aggregation despite light activation. These results are consistent with prior reports demonstrating that the extreme N-terminus is indispensable for membrane anchoring, with the first 25 amino acids, particularly the first 7, being critical for this interaction^29,68–71^. In line with this, PD-linked mutations that weaken helix-lipid interactions, namely A30P and G51D, similarly impaired recruitment and substantially reduced aggregation, whereas H50Q and A53T had little effect, and E46K further enhanced aggregation. These responses mirror biophysical studies showing that membrane-averse mutants such as A30P and G51D^47,48,72–75^ severely compromise membrane binding and consequently diminish primary nucleation. Yet, both A30P and G51D remain toxic and are linked to familial PD, suggesting that even a limited pool of α-SYN aggregates may be sufficient to elicit neurotoxicity despite reduced aggregation efficiency.

Of note, interaction with lysosomes does not appear to represent a canonical degradative process. Pharmacological inhibition of autophagy does not alter LIPA-α-SYN total protein levels, and high-resolution imaging demonstrates that aggregates remain attached to the lysosomal membrane without entering the lumen. This observation aligns with prior studies showing that dopamine-induced α-SYN oligomers, unlike monomeric or dimeric α-SYN, remain associated with the lysosomal membrane and escape chaperone-mediated autophagy^76–79^. Combined, these data suggest that lysosomes may play an active role in promoting α-SYN aggregation, beyond their classical function in protein quality control.

Although the selectivity of α-SYN for the lysosomal membrane can be partly explained by membrane biophysical properties, lipid composition (especially with the specific presence of bis(monoacylglycero)phosphate (BMP), a unique lysosomal anionic lipid of the inner leaflet, that accumulates in the brain^80^), and the high intrinsic propensity of the α-SYN N-terminus for membrane binding, these factors alone cannot fully account for the rapid and marker binding affinity of α-SYN aggregates to lysosomal surfaces. Upon light stimulation, α-SYN assemblies form directly on the lysosomal membrane within seconds, a timescale too fast to be explained by *de novo* anionic lipid remodeling, typically occurring over several minutes^81^. An alternative explanation is that α-SYN oligomers rapidly engage protein adaptors or scaffold complexes localized at pre-existing lysosomal microdomains, which tether and stabilize nascent assemblies. Supporting this hypothesis, our previous proteomic analysis revealed that early α-SYN oligomers interact closely with membrane-associated proteins^82^, suggesting these proteins may serve as adaptors facilitating their lysosomal localization. Among these proteins, WDR44 emerges as a strong candidate for anchoring and stabilizing lysosome-bound α-SYN oligomers. WDR44 selectively interacts with both α-SYN and lysosomes under aggregation-prone conditions, but not with monomeric α-SYN, consistent with an aggregation-dependent tripartite interaction that facilitates the conformational conversion of α-SYN into oligomers. Moreover, downregulation of WDR44 markedly reduced de novo α-SYN aggregation, whereas its overexpression enhanced α-SYN aggregation and inclusion formation, suggesting an important regulatory role for this protein in the α-SYN aggregation process. This protein, localized in the cytosol, play the role of a scaffolder that can efficiently clamp membranes, binding tubular endosomes (through the polyproline helix), recycling endosomes (Rab11BD, Rab11-binding domain) and the endoplasmic reticulum (ER) (FFAT domain)^49,50,83^. On the other hand, WDR44, as a member of the WD40-repeat family, is structured with a seven-bladed β-propeller that may act as a multivalent reader of short linear motifs and/or ubiquitylated proteins, accounting for its affinity for nascent α-SYN oligomers that are raising the valency^84,85^. Thus, α-SYN oligomers are multiplying key docking determinants for WDR44 in a single lysosomal membrane that may favor the binding of the WD40 β-propeller domains^86,87^. WDR44 may function as a sensor for α-SYN oligomers via its β-propeller domains, while recruiting multiple membranes to the vicinity of the aggregates through its Rab11-binding, FFAT, and polyproline helix domains^49,50,53,83–86^. We acknowledge, however, that WDR44 might not act alone, and may instead function in concert with additional proteins that we previously identified in our proteomic screen^23^, especially endolysosomal proteins that have already been directly or indirectly linked to PD such as VPS13C^88^, RABEP1^89^, COPB1^89^, BLTP3A^90,91^, HOOK1^92^ or IGF2R^93,94^, giving insights as to why WDR44 knockdown does not completely abolishing α-SYN aggregation. Among those proteins, VPS13C and BLPT3A, two lipid transporters associated with membranes, might be of high interest as WDR44 have been shown to directly interact with such proteins^83,95^.

Such scaffolding would hinder lysosome mobility, promote cargo retention, and concentrate anionic/curved membrane, aligning with the organelle-rich ultrastructure of LBs and our ultrastructural LIPA data^20,44,59,96^. The accumulation of the protein in the vicinity of the LBs, as well as the increase of WDR44 protein levels in PD and PDD brains, is relevant to the disease and in close alignment with large-scale label-based proteomics of LB dementias^58^. While not reported in the main highlights, WDR44 was specifically enriched in PD and PDD Brains, but not dementia with LB and Alzheimer’s disease, in multiple cohorts from UPenn (354 patients), Emory (44 patients) and Religious Orders Study or Rush Memory and Aging Project (ROSMAP, 103 patients out of the 610 retained) underscoring its association with α-SYN pathology in motor regions^58^. The functional consequences of α-SYN/WDR44/Lysosomes coupling are deleterious. Our data highlight that WDR44 overexpression exacerbated neuronal dysfunction in iPSC-derived human neurons through reduced axonal arborization and lysosomal alkalinization, linking WDR44/α-SYN complex to lysosomal failure and cell death. In line with our work, several reports point to lysosomal dysfunction as a core driver of PD pathogenesis and neuronal loss^97^. These deficits are also convergent with our previous findings, where we reported clustering of dysmorphic organelles, including lysosomes in the vicinity of LIPA-α-SYN aggregates at later time points and have observed nigral degeneration *in vivo*^20^. Notably, we also have previously detected BLTP3A at the interaction site of α-SYN/WDR44 complexes^23^. BLTP3A has recently been identified as a damage-responsive element that re-associates with injured lysosomes via an ATG8-interacting LIR motif consistent with a localized lysosomal stress response at these sites^95^. We also have observed *in vivo* that α-SYN aggregates associate with abundant multilamellar bodies, a hallmark of lysosomal lipid dyshomeostasis, aligning with ultrastructural studies that place vesicles and damaged organelles at the core of Lewy pathology^59^ and correlated with GBA1-induced PD^98,99^ or in other lysosomal storage disorders like Niemman Pick disease^100,101^.

In conclusion, our study uncovers a critical mechanistic step in the pathogenesis of PD, demonstrating that the initiation of α-SYN aggregation depends on its interaction with the lysosomal membrane and involves the protein WDR44. Notably, WDR44 is abnormally accumulated in the brains of PD patients and related synucleinopathies, where it correlates with the pathological α-SYN burden and colocalizes with Lewy bodies. This abnormal accumulation not only implicates WDR44 in the early stages of α-SYN aggregation but may also represent a novel biomarker for PD and related disorders. Together, these findings establish WDR44 as a promising therapeutic target and suggest that modulating its interaction with α-SYN could provide a novel strategy to prevent or slow the formation of pathological aggregates, thereby opening new avenues for disease-modifying interventions in PD and other synucleinopathies.

## Statistics

Data are reported as mean ± standard error of the mean (SEM) unless stated otherwise. Statistical comparisons between two groups were performed by using appropriate tests as described in each figure legend, ensuring sample normality and equal distribution of standard deviation. A p-value of less than 0.05 was considered statistically significant. Statistical analysis and graphs were generated using GraphPad Prism software (v.10). All *in vitro* and *in vivo* experiments were independently replicated at least three times; additional repeats for key experiments are noted in the figure legends. All replication attempts yielded consistent results. Representatives immunoblot images and microscopy images reflect at least three independent experiments, unless otherwise specified in the figure legends.

## Supporting information

Suppl vSideo 1

## Materials and Methods

### Plasmid cloning

Tetracycline-inducible pLVX 3K-α-SYN-YFP (gift from Dr. Ulf Dettmer) was previously characterized and described^24,26,102^. pcDNA and pAAV CMV templates of Cry2olig-mCherry (LIPA-EMPTY), α-SYN-Cry2olig-mCherry (LIPA-α-SYN) and TDP-43-Cry2olig-mCherry (LIPA-TDP-43) were generated and described in our previous work^20,23^.

Mutations of α-SYN within the LIPA-α-SYN backbone were performed using overlapping mutagenic primers to amplify the whole backbone, incorporating the mutations. Forward primers were designed as follows: 5’-GCA GAA GCA CCA GGA AAG AC-3’ for A30P, 5’-CAA AAC CAA GAA GGG AGT GGT G-3’ for E46K, 5’-GTG GTG CAG GGT GTG GCA A-3’ for H50Q, 5’-GTG GTG CAT GAT GTG GCA AC-3’ for G51D and 5’-GCA TGG TGT GAC AAC AGT GG-3’ for A53T. Reverse primers were designed as follows: 5’-GTC TTT CCT GGT GCT TCT GC-3’ for A30P, 5’-CAC CAC TCC CTT CTT GGT TTT G-3’ for E46K, 5’-TTG CCA CAC CCT GCA CCA C-3’ for H50Q and 5’-GTT GCC ACA TCA TGC ACC AC-3’ for A53T.

N-terminal truncations of α-SYN within the LIPA-α-SYN backbone were performed using mutagenic primers amplifying the region of interest, adding a Methionine whenever necessary and an AfeI cleaving site upstream of the Methionine. Forward primers were designed as follows: 5’-GGG GAC AGC GCT ATG AAA GGA CTT TCA AAG GC-3’ for α-SYN^Δ2-4^, 5’-GGG GAC AGC GCT ATG GCC AAG GAG GGA GTT GTG GC-3’ for α-SYN^Δ2–10^, 5’-GGG GAC AGC GCT ATG GCT GAG AAA ACC AAA CAG GG-3’ for α-SYN^Δ2–18^. A generic reverse primer was designed as follows to amplify the rest of the insert, covering up to a downstream MfeI cleaving site: 5’-CAA GTT AAC AAC AAC AAT TGC ATT CAT TTT ATG TTT CAG G-3’. Non-mutated insert was removed from its backbone using AfeI (NEB, cat. R0652S) and MfeI-HF (NEB, cat. R3589S), and the backbone lacking the insert was purified using PureLink™ Quick Gel Extraction and PCR Purification kit (Thermofisher, Cat. K220001). Following the same method, PCR products containing the amplified inserts (with truncations) were purified, digested with AfeI and MfeI and then ligated using T4 DNA ligase (NEB, Cat. M0202T), following a ratio of 3:1 (insert:vector). The ligated material was finally transformed in DH5-α competent cells (NEB, Cat. C2987H) and amplified using GenElute™ HP Plasmid Maxiprep Kit (Sigma, Cat. NA0310-1KT). C-terminal truncations of α-SYN within the LIPA-α-SYN backbone was performed following the same general method with adjustments. A generic forward primer was designed as follows to amplify the start of α-SYN, amplifying an AfeI cleaving site upstream of the insert: 5’-CCG GAC TCA GAT CTC GAG GAC AGT GTG G-3’. Mutagenic primers were designed to amplify α-SYN up to the specific amino acids, inserting a stop codon and a KpnI cleaving site downstream. Reverse primers were designed as follows: 5’-CCC CGG TAC CAT TCT TGC CCA ACT GGT CC-3’ for α-SYN^Δ104–140^, 5’-CCC CGG TAC CTT CCA GAA TTC CTT CC-3’ for α-SYN^Δ115–140^, 5’-CCC CGG TAC CAG GAT CCA CAG GCA TAT CTT CC-3’ for α-SYN^Δ121–140^ and 5’-CCC CGG TAC CAT CTT GAT ACC CTT CCT CAG AAG GCA TTT C-3’ for α-SYN^Δ136–140^.

PiggyBac vector containing LIPA-α-SYN was obtained from the vector PiggyBac-rtTA-sfGFP-IRES-NGN2-puro (Addgene, cat. 209079)^103^ into which the α-SYN-Cry2olig-mCherry insert was cloned. PiggyBac vector containing the WDR44 gene was introduced separately using a second inducible PiggyBac-compatible backbone PiggyBac-TA-ERN (Addgene, cat. 80474)^104^ that allows for a high overexpression while maintaining a separate antibiotic selection. The cloning strategies were achieved via standard molecular cloning methods, Gateway® BP and LR Clonase™ reactions, following the manufacturer’s instructions (Thermo Fisher Scientific, cat. 11789020 and 11791020). Plasmids integrity have been validated through PCR and complete sequencing of the inserts for all the detailed cloning experiments.

pGCS2-hSyn-Cry2olig-mCherry and pGCS2-Cry2olig-mCherry were created using the Gateway® BP and LR Clonase™ reactions, following the manufacturer’s instructions as described earlier, using pDEST pGCS2 empty backbone as a destination vector (Addgene, cat. 85731).

### Human tissue

#### Canada – Saskatchewan brain deposit on movement disorders

Brain samples were obtained from the Saskatchewan Movement Disorders Program (SMDP) where clinical data were collected from patients, allowing for a longitudinal follow-up^57,105^. Consent for brain autopsy and use of brain tissue for research was approved by the hospital ethics committee and by the University of Saskatchewan Ethics Board^57,105^. Clinical evaluations were accomplished by one of two movement disorders neurologists (AHR, AR), and the post-mortem diagnosis was done for each patient by a Canadian certified neuropathologist^57,106^. Data such as sex, age at onset, duration of disease, brain weight, and presence of Lewy Bodies were collected. The Hoehn & Yahr (H & Y) scale, being the most widely used and accepted staging system, was initially used to measure the global severity, followed by the use of the Unified Parkinson’s Disease Rating Scale (UPDRS) and modified H & Y scales (stages 2-3.5: n=10; stages 4-5: n=14)^105,107^. The control individuals were younger than the patients with PD. In addition, the PD group had a higher proportion of men compared to the control group, which is consistent with evidence reporting that the risk of developing PD is two times higher in men compared to women^108^. Autopsies were performed within 24 hrs after death, and the collected brains were separated into two: one half of the brain, which was fixed in formalin, was used for histologic and diagnostic studies of the midbrain, while the other half was frozen at-80°C and cut along the frontal plane to obtain 2-3 mm thick slices. Slices corresponding to the parietal cortex, the putamen, the SN (coronal plane) and the cerebellar cortex (axial plane) were used to extract tissues for these regions (∼100 mg). Coronal slices containing the SN were cryostat-sectioned (20 µm), thaw-mounted onto SuperFrostPlus slides (75X50 mm), desiccated overnight at 4°C, and stored at - 80°C until assayed, as described^109^. In addition, the SN were dissected on cryostat sections and 5 x 50 µm sections were harvested to obtain approximately 30 mg of frozen sample and stored at - 80°C. Tissue pH was measured as an indication of tissue quality^109,110^.

#### Canada – Quebec CERVO brain deposit

Tissue samples were also obtained from the human brain bank at the CERVO Brain Research Centre. Informed consent was required before donation. The Ethics Committees at Université Laval and CIUSSS de la Capitale Nationale approved the brain collection procedures and the storing and handling of postmortem human brain material. Analyses were performed in accordance with the Code of Ethics of the World Medical Association (Declaration of Helsinki). Immunofluorescence assays were performed on sections from the brains of 5 patients who suffered from PD (PD: mean age 73 ± 5 years and postmortem delay (PMD) ranging from 2.5 to 18 hrs, **Table II**), with no evidence of other cognitive, psychiatrice or neurological disorders. Brains sections from 5 additional patients with PD and dementia (PDD: mean age 79 ± 8 years, PMD ranging from 4 to 58 hrs) were also used. Movement disorder neurologists performed all clinical diagnoses and collected clinical data such as age at onset, duration and severity of the disease and dopamine medication as calculated with levodopa equivalence dose^111^. Clinical diagnoses were confirmed following a detailed pathologic examination performed by a certified neuropathologist according to the Lewy pathology consensus criteria^112–114^. Only patients who suffered from PDD demonstrated a diffuse neocortical form of Lewy body type pathology^112^.

#### France – Bordeaux brain deposit

Formalin-fixed and paraffin-embedded human brain samples from PD patients were obtained from the repository of the University Hospital Bordeaux (CRB-BBS). Written informed consent was obtained for the collection of the brain and the use of clinical and post-mortem data from all subjects or their legal representatives and mandatory regulatory approval for post-mortem brain tissue use was obtained (MESR AC-2024-6715, DC-2024-6714).

### Cell Maintenance and differentiation

#### Immortalized cell lines

HEK293T (ATCC, cat. CRL-3216) and COS-7 cells were maintained in high-glucose Dulbecco’s modified Eagle’s medium (DMEM) (Sigma-Aldrich, cat. D5706) supplemented with 10% fetal bovine serum (FBS) (Sigma-Aldrich, cat. F1051) and 1% penicillin/streptomycin (ThermoFisher, cat. 15-140-122) at 37°C and 5% CO_2_.

#### Human induced pluripotent stem cells (hiPSCs)

The hiPSC line used in this study was derived from the parental AIW002-02 hiPSC line, which was described previously and analyzed for multiple quality control parameters, including pluripotency by immunofluorescence, trilineage differentiation potential and genomic integrity using the hiPSC Genetic Analysis Kit (Stemcell Technologies, cat. 07550). 3X *SNCA* hiPSC line (originally named AST23 or ND34391*H (RRID: CVCL_F202)) were obtained as a gift from Dr. Thonas Durcan The hiPSCs used in the study was approved by the CHU de Quebec Research Center (#2022-6079). The hiPSCs line was cultured and maintained under standard conditions, as previously described^34,35^.

Cells were grown in mTeSR plus medium (STEMCELL Technologies, cat. 100-0276) on Matrigel-coated plates (Corning, cat. 354277) and passaged using 0.5M EDTA-based dissociation reagent. Cells were then differentiated into dopaminergic neurons (iDA) as previously described^34,35^ using STEMdiff^TM^ differentiation and maturation kits (STEMCELL Technologies, cat. 08520 and 08530).

### Induction of protein aggregation

HEK293T and COS-7 cells were seeded on poly-L-lysine coated glass coverslips to reach 70% confluency the next day. 12 hrs post plating, cells were transiently transfected with LIPA-α-SYN, LIPA-EMPTY, LIPA-TDP-43 or pLVX 3K-α-SYN-YFP using calcium phosphate (HEK293T) as previously described^20,21^ or following the Lipofectamine® 2000 protocol (COS-7). For the LIPA-constructs, 16 hrs post-transfection, cells were exposed to blue light (λ = 456 nm) using UHP-T-DI-LED series Ultra High-Power LEDs (Prizmatix, Southfield, MI, USA), at the intensity of 0.30 mW/mm^2^ measured using LPM-100 light power meter (Amuza, San Diego, CA, USA). For the 3K-α-SYN-YFP, 16 hrs post-transfection, the cell culture medium was replaced with medium containing doxycycline at 1 µg/ml, allowing the induction of expression of the construct. Doxycycline-containing medium was replaced every day to spike fresh doxycycline, and cells were fixed at different time points (see figures). For colocalization assays, iDA neurons^34,35^ were transiently transfected after 18 days *in vitro* (DIV) using Lipofectamine®2000 with a 1:3 (DNA: lipofectamine) ratio. Briefly, neuronal conditioned medium was collected from each well and kept on ice. Opti-MEM medium was added to the culture. In parallel, DNA and lipofection reagent were mixed to form DNA-liposome complexes and incubated for 20 min, then added to the cells for 4 h. After incubation, the culture medium was replaced with a 1:1 (v/v) mixture of the previously saved conditioned medium and fresh maturation medium. Thirty-six hours post-transfection, neuronal cultures were used for the experiments and further exposed to blue light following the same parameters described earlier, using a reduced blue light power (0.15 mW/mm^2^) to minimize phototoxicity making sure that cells not transduced with LIPA were not affected. For toxicity and population-based assays, stable iPSC-lines were generated using PiggyBac transposons. We used a PiggyBac transposon-based delivery system to stably integrate the LIPA module and WDR44 gene in our NGN2-hiPSC line (see cloning paragraph). iPSCs were co-transfected with the PiggyBac vector(s) containing either one or the other insert (LIPA-α-SYN and/or WDR44), and a hyperactive PiggyBac transposase plasmid (pCMV-hyPBase) (a gift from the Canadian Neurophotonics Platform-CERVO), using Mirus TransIT®-LT1 Transfection Reagent (Mirus, cat. MIR 2304) as previously described^23^. Following antibiotic selection (puromycin at 1 µg/ml (NGN2), blasticidin at 50 µg/ml (LIPA-α-SYN), G418 at 75 µg/ml (WDR44)), integration was confirmed via protein expression levels. Following any genomic integration, the modified hiPSC lines were evaluated for maintained pluripotency and differentiation potential via immunocytochemistry for key lineage-specific markers. Evaluation of the protein expression was assessed at DIV 10 through Western blot to ensure WDR44 overexpression and equal LIPA-α-SYN expression in the cells overexpressing WDR44 as compared to the endogenous-WDR44 line.

### Transduction of shRNA to generate stable cell lines with reduced protein expression

To generate stable HEK293T with reduced protein expression, we used the shRNA system pLKO.1 vectors that were picked from glycerol stocks of the publicly available The RNAi Consortium (TRC) libraries (Sigma-Aldrich, MO). Plasmids were recovered from glycerol stocks, expanded, pooled, and purified by maxiprep (gift from Dr. Stephane Gobeil). At least two shRNA vectors were tested per protein of interest: WDR44 (TRC 062495, TRC 062494), IGF2R (TRC 060255, TRC 060257), BLTP3A (TRC 064774, TRC 064773), Scramble (TRC scramble, addgene cat. 1864). Cells were transduced with lentiviral particles purified for each construct and further selected with puromycin 1 µg/ml. After confirming robust knockdown at both mRNA and protein levels, we selected the constructs producing the greatest reduction for subsequent experiments.

### Animals

#### Thy1-α-SYN mice

For this study, heterozygous adult male mice over-expressing human WT α-SYN driven by the murine Thy1 promoter (Thy1-α-Syn mice, line 61) on a C57BL/6 background and littermate controls were used^115^. The mice were subsequently maintained as an in-house colony at the Centre de Recherche du CHU de Québec–Université Laval (CHUQ-UL) and were housed four per cage in ventilated cages with standard rodent bedding in a temperature and light-controlled room (22 °C, 12 hrs cycle) with ad libitum access to food and water. Animals were sacrificed at 8 months of age, and brains were sliced as previously described^116^. All experiments were performed in accordance with the Canadian Guide for the Care and Use of Laboratory Animals, and all procedures were approved by the Institutional Policy of the CHUQ-UL (protocol 2017-128 to FC).

#### LIPA-α-SYN AAV mice with optogenetic stimulation

Three-month-old C57/BL6 mice were obtained from Charles River Laboratories and habituated for 7 days before any handling. Mice were housed and fed as described earlier, and all animal experiments were approved by the Animal Welfare Committee of Université Laval in accordance with the Canadian Council on Animal Care policy, protocol 2020–641 to AO. Optogenetic implants surgery, LIPA-α-SYN AAV particles delivery and blue light stimulation was performed as previously described^20^. Briefly, fifteen days after the surgery where implants and AAV-delivery have been performed, i*n vivo* blue light stimulation sessions were performed every other day using the wireless optogenetic devices (Eicom/Amuza). Right before each session, a battery (Eicom/Amuza) was directly connected to the implant. Stimulation was performed for 1 hr using a pulse generator (Eicom/Amuza) at the power of 1.76 mW/mm^2^ (measured at the tip of the optical fiber) and a pulse of 10 ms at 20 Hz. Animals were sacrificed after 7 days of stimulation.

#### Zebrafish micro-injection and aggregation assays

Zebrafish experiments were performed in accordance with protocols (ASP#20-416) approved by the animal care & use committee of the National Cancer Institute at Frederick in compliance with the Association for Assessment and Accreditation of Laboratory Animal Care (AAALAC) guidelines. AB zebrafish were maintained under a 14 h light/10 h dark cycle at 28 °C. One-cell stage embryo were co-microinjected with 250 pg/nL of pCSG2-hSyn-Cry2olig-mCherry and either 250 µM of a standard control morpholino (Ctrl MO 5’-CCTCTTACCTCAGTTACAATTTAT-3’) or 250 µM of the translation blocking *wdr44*MO (5’-GCTGCATACAGAGGCCGCCGCCACT-3’)^54^ both obtained from Gene Tools. Capped mRNAs were *in vitro* transcribed using the mMessage mMachine SP6 Transcription Kit procedure (ThermoFisher, cat. AM1340). For aggregation assays, embryos were maintained in E3 media in the dark post-injection at 28 °C. At 24 hrs post fertilization (hpf), embryos were placed in a Nunc™ Lab-Tek™ II Chambered Coverglass (ThermoFisher Scientific, cat. 155360) with E3 media. Cells on the surface of the yolk of live embryos were exposed to the 488nm laser at 37 mW/mm^2^ (to induce the activation of the cry2 system) and Z-stack images with an axial step size of 0.3µm and range of 6µm were captured at t0 minute and every 5 sec for 10 min using the 63x 1.3 NA Zeiss oil objective on Marianas spinning disc confocal microscope (SDCM) (Intelligent Imaging Innovations) equipped with a sCMOS (Hamamatsu ORCA Flash4) camera. The percentage of mCherry cells with positive aggregates per embryo (cells with a minimum of one aggregate were counted) is calculated as the number of cells with mCherry-positive aggregates/total number of mCherry-positive cells.

### Immunofluorescence assays

#### In vitro

Immortalized cell lines (HEK293T or COS-7), plated on coverslips, were fixed with 4% PFA, 3% Sucrose for 15 min at RT and further washed three times 5 min with Phosphate Buffer Saline (PBS). Cells were permeabilized for 5 min with 0.1% Triton and washed once time with PBS. Blocking was performed with PBS-0.1% Saponin 5% Normal Goat Serum (NGS) and 2% Bovine Serum Albumin (BSA) for 1 hr at RT. Cells were then incubated with primary antibodies (table III): LAMP1 (Abcam, cat. 24170), LAMP2 (Invitrogen, cat. MA1-205) for 1 hr and washed three times for 5 min with PBS before incubation with matching secondary antibodies (see table I). After three washes for 5 min with PBS, cells were counterstained with DAPI and coverslips mounted with ProLong diamond antifade (Invitrogen, cat. P36965). For iPSC-derived neurons, the same pipeline was followed with minor adaptations: initial permeabilization was performed with 0.3% Triton for 30 seconds, and the blocking buffer was composed of PBS-0.1% Triton, 5% NGS, 2% BSA. Specific primary antibodies were used to assess neuronal phenotype (table III): TH (Phosphosolutions, cat. 2025-THRAB), TH (Aves Labs, cat. TYH-0020).

**Table III.**
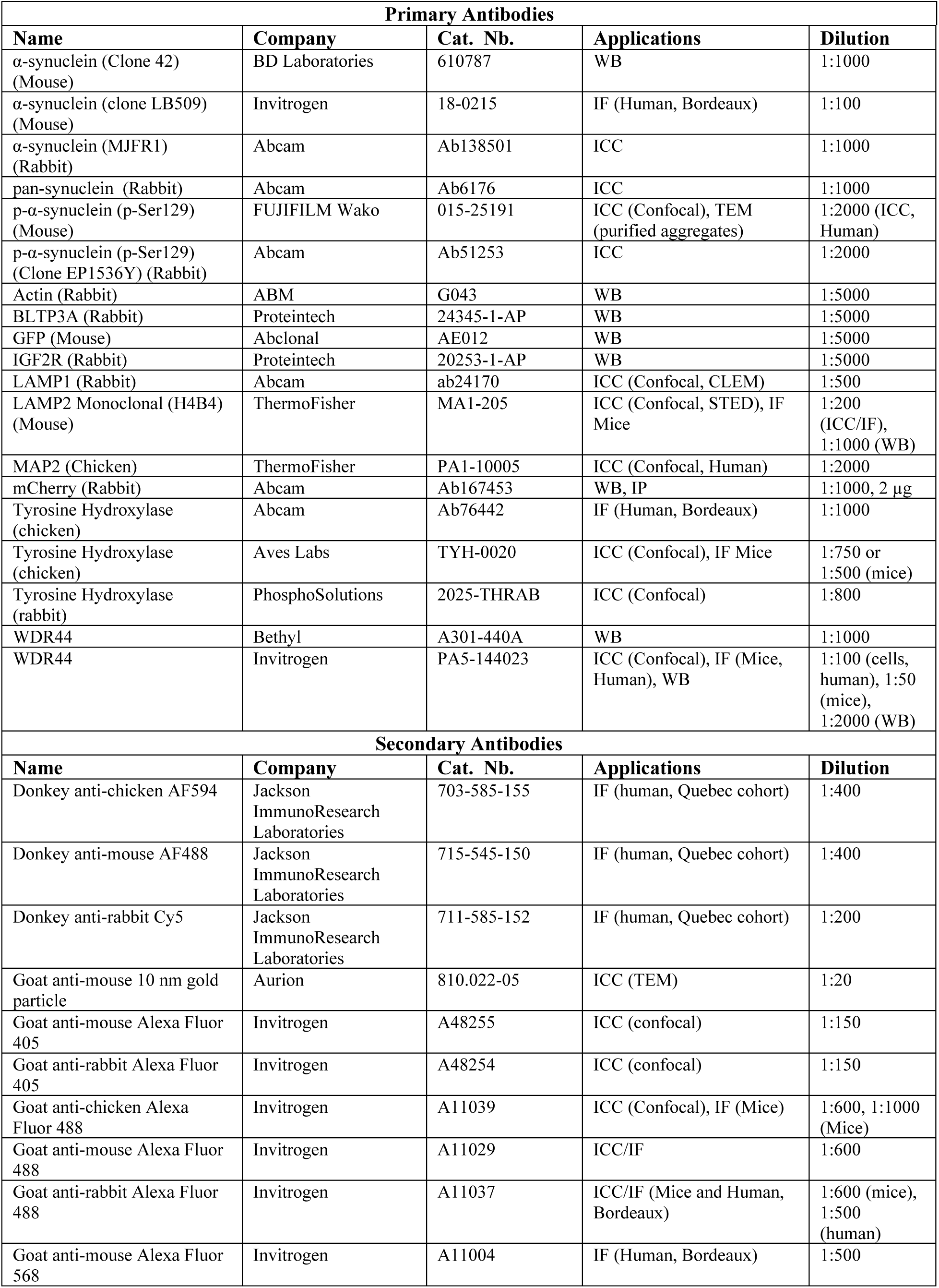
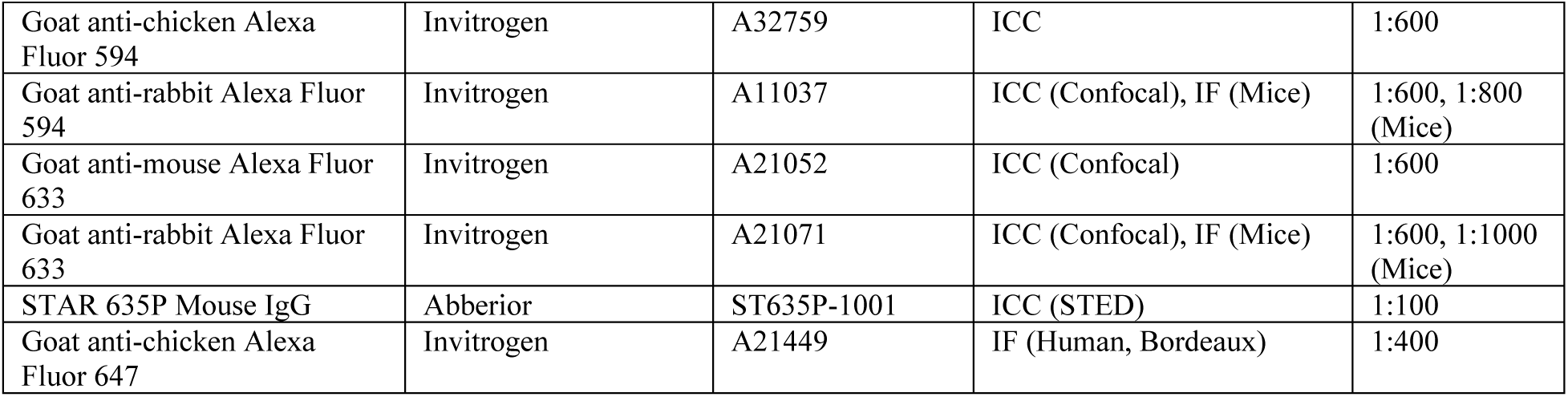
Antibody list, sources, applications and dilution factors.

***In vivo***. Thy1-α-Syn free-floating brain sections of the substantia nigra were washed with KPBS and incubated with citrate buffer (pH 6.2) for 30 min at 85 °C for antigen retrieval. Following three KPBS washes (8 min each), the brain slices were blocked for 30 min at RT in a KPBS solution containing 4% NGS (Wisent Bioproducts, cat. 053-150), 0.1% Triton X-100, 1% BSA. The slices were then incubated overnight at 4 °C with anti-WDR44 and anti-TH antibodies (see Table III). The following day, the substantia nigra sections were washed three times (10 min each) with KPBS and subsequently incubated with secondary antibodies (see table I) for 2.5 hrs at RT. Brain sections were then washed three times for 10 min in KPBS, followed by an incubation with 4′6-diamidino-2-phenylindole (DAPI) nuclear stain (Invitrogen, cat. D35571, 1:2000) diluted in KPBS for 7 min. Finally, sections were washed with KPBS (three times, 5 minutes each) and mounted on slides (Fisher Scientific, cat. 12-550-15) using Fluoromount-G (Invitrogen, cat. 00-4958-02). LIPA-α-SYN free-floating brain sections of the substantia nigra were stained following the same protocol, without antigen retrieval and using primary antibodies WDR44, TH or LAMP2 (see table I).

#### Human brain samples – Quebec CERVO brain deposit

Brains were sliced in half along the midline and hemi-brains were cut into 2-cm-thick slabs along the coronal plane. The slabs were fixed by immersion in 4% paraformaldehyde at 4°C for 3 days and stored at 4°C in a 0.1M PBS, pH 7.4 solution containing 15% sucrose and 0.1% sodium azide. Slabs containing the SN were cut with a freezing microtome into 50 µm-thick transverse sections that were serially collected and stored at 20 °C in a solution containing glycerol and ethanediol until immunostaining. One section was sampled from each brain using the Human Brain Atlas as a reference^117^. The 50 μm-thick free-floating sections were washed 3 times in PBS, followed by an incubation at 100 °C for 1 hr in an antigen retrieval solution (Sodium citrate 10mM, 0,05% Tween 20, pH 6.0). Sections were washed three times in PBS, blocked for 2 hrs in PBS containing 2% species-appropriate normal serum and 0.1% Triton X-100, then incubated overnight at 4 °C with primary antibodies (Table III) diluted in the same blocking buffer. The following primary antibodies were used: chicken anti-MAP2 (Invitrogen, cat. PA1-10005), rabbit anti-WDR44 (Invitrogen, cat. PA5-144023), and mouse anti-phosphorylated α-synuclein (FUJIFILM Wako, cat. 015-25191). On the following day, sections were rinsed three times in PBS, incubated with the corresponding secondary antibodies for 2 hrs (Table III), washed three times in PBS, mounted on positively charged slides, air-dried, quenched with Autofluorescence Eliminator Reagent (MilliporeSigma), and mounted with Dako Fluorescence Mounting Medium (Agilent, cat. S3023).

#### Human brain samples – Bordeaux brain deposit

To detect the presence of WDR44 protein in Lewy bodies, sequential immunofluorescence labelling was performed on 4-µm coronal paraffin-embedded sections of the substantia nigra using an anti-WDR44 antibody (rabbit polyclonal, Invitrogen, cat. PA5-144023) and an anti-tyrosine hydroxylase (TH) (chicken polyclonal, Abcam, cat. Ab76442), a marker of DA neurons, coupled to the anti-α-synuclein antibody (mouse monoclonal clone LB509, Invitrogen cat. 18-0215). First, tissue sections were deparaffinized in xylene and rehydrated through a graded series of ethanol from 100 to 70% before undergoing antigen retrieval by pressure cooking (TintoRetriever Pressure Cooker) at 100 °C–112 °C for 5 minutes in EDTA buffer pH 9.0 (Agilent-DAKO, cat. S236784-2) followed by 5 min in citrate buffer pH 6.0 (Agilent-DAKO, cat. S169984-2). After cooling, sections were washed in Tris-buffered saline containing 0.3% Triton X-100 (TBS 0.1M - 0.3% Triton X-100) and blocked with 5% normal goat serum and 1% BSA in the same buffer for 1 hr at RT. Sections were then incubated overnight at 4°C with the anti-WDR44 antibody revealed with goat anti-rabbit Alexa fluor 488 (1:500). Subsequently, sections were exposed overnight at 4 °C to a mixture of the anti-TH and anti-α-synuclein primary antibodies. Incubation with secondary antibodies was performed sequentially using goat anti-mouse Alexa Fluor 568 (1:500) for α-synuclein and goat anti-chicken Alexa 647 (1:400) for TH. To reduce lipofuscin autofluorescence, slides were incubated for 10 min in 0.1% Sudan Black B (Sigma-Aldrich, cat. 199664) in 70% ethanol. Nuclei were counterstained with 4’,6-diamidino-2-phenylindole (DAPI, Sigma-Aldrich, cat. D9542). After thorough washing in TBS, slides were mounted in fluorescence mounting medium (Agilent DAKO, cat. S-3023). For co-localization analysis, images were taken at 40x magnification using a computerized image analysis system (Morphostrider, Explora Nova) connected to a Zeiss Axioplan 2 epifluorescence microscope.

### Confocal Microscopy

#### Live-cell aggregation induction and analysis

For live-cell assessment of the interaction between LIPA-α-SYN and the endolysosomal system, COS-7 cells were plated at 70% confluency in poly-D-lysine-coated glass-bottom 35-mm dish (Mattek, cat. P35GC-1.5-14-C). The next day, cells were co-transfected (1 µg LIPA-α-SYN with 0.75 µg OTC-GFP (gift from Dr. Étienne Hébert-chatelain), 0.5 µg GFP-RAB5B (Addgene, cat. 61802), 0.5 µg GFP-RAB7 (Addgene, cat. 12605) or LAMP1-mGFP (Addgene, cat. 34831)) in Opti-MEM™ (ThermoFisher, cat. 31985070) using Lipofectamine® 2000 (Thermofisher, cat. 11668027) with a 1:3 (DNA: lipofectamine) ratio. After 16 hrs of transfection, the culture medium was replaced with FluoroBrite™ DMEM (ThermoFisher, cat. A1896701) supplemented with 10% FBS and GlutaMAX™ (100X dilution, ThermoFisher, cat. 35050061). The cells were then placed in a stage-top incubator (Tokai Hit) on the microscope, with CO maintained at 5% and temperature at 37 °C. After 30 min of stabilization, time-lapse imaging was initiated using an inverted Leica TCS SP8 STED 3X microscope in confocal xyzt mode, equipped with an HC PL APO CS2 63x/1.40 OIL objective, 3.5x numerical zoom, 1248×1248 pixel resolution, unidirectional resonant scanning at 8000 Hz with 4-line averaging. Both wavelengths (GFP and mCherry) were acquired simultaneously using a white light laser (WLL) set at 70% input power using two individual Hybrid detectors (HyD) gated for their respective excitation lasers to maximize the temporal resolution (which also permitted continuous 488-nm laser stimulation of CRY2olig), and z-stacks were acquired with 0.5 µm steps (8 µm total thickness) to capture all images for both sequences within 10.249 sec. Each stack was followed by continuous imaging for up to 90 min. Fluorescence was recorded using HyD detectors with the following settings: GFP (15% laser power at 488 nm, fluorescence collected at 494-530 nm, gain 200V, 0.30-6.00 ns time gating) and mCherry (5% laser power at 587 nm, fluorescence collected at 605-750 nm, gain 150V, 0.30-6.00 ns time gating). For the co-localization analysis, manual quantifications were performed on z-stack projections at key timepoints of continuous imaging (10s, 30s, 1mn, 1mn30s, 5mn, 15mn, 30mn, 60mn and 90mn) to assess the percentage of LIPA-α-SYN aggregates localized to the membrane of an organelle. Co-trafficking events were assessed using kymographs generated with the open-source software Fiji (https://imagej.net/software/fiji/). Briefly, a representative Region of interest (ROI) was defined at 5 and 30 min of live imaging; a line (width = 20) was placed to track particles across the intervening 25-min interval. The plugin KymographBuilder (https://imagej.net/plugins/kymograph-builder) was then applied to obtain the kymograph along the line described earlier. Analysis of the lysosomal network was performed on segmented images using weka segmentation in ImageJ as a built-in option from Trackmate 7^118^. Once organelles were segmented in the first 15 min and the last 15 min of live-cell imaging, individual lysosomes were tracked using Trackmate and lysosomal traveling distance was analyzed alongside the total individually tracked lysosomes. Color-coded individual tracks showing the distance traveled were also generated through Trackmate^118^.

#### Neuronal lysosomal dysfunction analysis in living cells

To assess the lysosomal pH and health of neurons undergoing LIPA-α-SYN aggregation and/or WDR44 overexpression, hiPSC-derived neurons (WT, LIPA-α-SYN, LIPA-α-SYN+WDR44) were exposed or not to blue light as stated in the figure. After each time point, cells were incubated with fresh medium containing LysoTracker deep red at 75 nM (Invitrogen, cat. L12492) and NeuroFluor NeuO at 1:200 (STEMCELL, cat. 01801) for 30 min in the dark. After 30 min, the media was replaced with fresh culture media and cells were directly imaged using an inverted Leica TCS SP8 STED 3X as previously described, with the following adaptations: 2x numerical zoom, 1024×1024 pixel resolution, bidirectional point scanning at 600 Hz with 2-line averaging. Three wavelengths were acquired, two simultaneously (NeuO and LysoTracker) through two individual Hybrid detectors gated for their respective excitation and one after the first sequence (mCherry). Parameters were set as follows: NeuO (10% excitation at 488 nm, collecting emitted fluorescence between 494 and 561 nm with a time gating of 0.30-6.00 ns), Lysotracker (1% excitation at 645 nm, collecting emitted fluorescence between 656 and 799 nm with a time gating of 0.60-8.00 ns) and mCherry (7% excitation at 587 nm, collecting emitted fluorescence between 592 and 629 nm with a time gating of 0.50-8.00 ns). For each condition, five large fields were acquired. Images were thresholded in ImageJ, and LysoTracker-positive puncta within each neuronal soma ROI were counted manually.

#### Neurites network analysis

To assess the impact of lysosomal dysfunction on neuronal health, we used the same experimental conditions as described in the lysosomal dysfunction analysis, without the use of LysoTracker or NeuroFluor NeuO. After each time point of blue light (or dark), cells were fixed with 4% PFA, 3% Sucrose for 15 min and immunofluorescence was performed as described earlier, using chicken anti-MAP2, rabbit anti-pS129-α-SYN (table I) as primary antibodies. Images were acquired using an inverted Leica TCS SP8 STED 3X microscope in confocal xyzt mode, and neurite length, number of branches and Sholl analysis were done using Simple Neurite Tracer’s moniker (SNT) plugin in imageJ on MAP2 signal^119^.

#### Quantification of number of cells with aggregates and number of aggregates per cell

Using an inverted Leica TCS SP8 STED 3X microscope in confocal xyzt mode, equipped with an HC PL APO CS2 63x/1.40 OIL objective, 1.5x numerical zoom, large fields of view were taken (at least 30-50 cells per field) to assess the number of cells with aggregates as well as the number of aggregates per cell. The number of aggregates per cell was manually quantified on Z-stack maximum intensity projections. An aggregate was considered a unique object when its perimeter could be traced as a continuous closed line; counts were performed with the Cell Counter plugin from Fiji/ImageJ. For the percentage of cells bearing aggregates, the total number of mCherry cells was manually quantified using the same method, and a cell bearing at least one delineated aggregate was considered a cell bearing an aggregate and quantified. To obtain the final percentage, the counted cells were merged from 3 to 6 fields of view (50 to 150 total cells per biological replicate).

#### Quantification of aggregates localization to organellar membranes and the colocalization surface

Aggregates and organelles were segmented directly from the z-stack using the plugin Distance Analysis (DiAna)^120^ in the open-source software Fiji /ImageJ (https://imagej.nih.gov/ij/), as previously described^23^. To optimize segmentation, detection and Gaussian-based segmentation thresholds were calibrated on the mCherry channel according to signal intensity, then held constant within each experiment. Objects with areas <10 px (background) or >90,000 px (aberrant signal, dead cells/debris) were excluded. Organelles were segmented using the same approach with specific thresholds that were held constant for each experiment and organelle type. Two objects were scored as interacting when they overlapped by ≥5 pixels, and the colocalization overlap was computed by the plugin at the same time. For highly crowded fields, such as the neuronal soma, causing segmentation errors, manual quantification was used instead.

#### Intensity profiles to assess colocalization

To generate fluorescence intensity profiles displaying the spatial localization of the fluorophores of interest, a line was drawn over the ROI and the fluorescence intensity was measured using the Plot Profile option from Fiji.

#### Quantification of WDR44 fluorescence level in Thy1-α-SYN mice

Following immunofluorescent labeling for WDR44 and TH, as described above, confocal z-stacks were acquired in the *substantia nigra* sections using an AXI0 Imager Z.2 upright confocal microscope equipped with a PL APO 40X/1.40 objective, 2X numerical zoom, and a unidirectional laser scanning at 700 Hz. For the analysis, maximum intensity projections were performed from the z-stacks and WDR44 intensity for each soma was quantified using the integrated density, calculating the corrected total cell fluorescence (CTCF) [integrated density – (area*average mean background)]. Quantifications were performed on 24 cells for the Thy1-α-SYN group and 25 cells for WT group.

### Stimulated emission depletion (STED) microscopy

For STED microscopy, COS-7 cells were seeded at 70% confluency on 1.5H precision coverslips (Thorlabs, cat. CG15NH1) coated with 0.1 mg/ml of Poly-L-Lysine (ScienCell, cat. 0403). Cells were then transfected with the LIPA-α-SYN plasmid using Lipofectamine® 2000 as previously described. Cells were then fixed with 3% PFA, 0.1% Glutaraldehyde for 15 min, followed by immunostaining as previously described. The following imaging parameters^121^ were used: STED images were acquired using an inverted Leica TCS SP8 STED 3X microscope, equipped with a motorized stage and a tunable white-light laser (470-780 nm) as well as a 405-nm diode laser (Leica Biosystems, Concord, ON, Canada). Images were acquired by sequential scanning between stacks with a 590-nm laser (LIPA-α-SYN, mCherry) or 656-nm (LAMP2, Abberior STAR 635P). Depletion of the fluorescence emitted by the mCherry or Abberior STAR 635P was performed using a 775-nm pulsed laser set at 30% input power and further tuned at 5%. Excitation laser intensities and gain were modulated to optimize the signal-to-noise ratio without saturation using the range indicator QLUT Glow mod. Point laser scanning was performed using a bidirectional scanner set at 1000 Hz, with 4-line averages. For all the images, a 100x HC PL APO CS2/1.40 NA STED white (Leica, cat. 11506378) was used with an immersion oil Leica type F (refractive index: 1.5180, Leica, cat. 11513859) and a x9 numerical zoom was applied. The wavelength emitted by the fluorophores was collected using Hybrid detectors (HyD) in standard mod, maintaining the digital gain below 100V, and using the time-gating option (0.3-6.0 ns). The pixel size was set to 15 nm, the size of the pinhole and z-step were set to optimize resolution or oversample for further deconvolution (voxel size of 130 nm). Deconvolution was performed using Huygens Professional (Scientific Volume Imaging, Hilversum, the Netherlands) using a theoretical point spread function, manual settings for background intensity, default signal-to-noise ratio, and a CMLE algorithm. Color balance, contrast, and brightness were adjusted using the open-source software Fiji/ImageJ (https://imagej.nih.gov/ij/). 3D surface rendering was performed on deconvoluted images using Imaris software version 7.6.1 (Bitplane, Zurich, Switzerland).

### Correlative-Light Electron Microscopy

#### In vitro

COS-7 cells were cultured as previously described on 35 mm glass-bottom dishes etched with alphanumeric coordinate grids (MatTek Corporation, P35G-1.5-14) to facilitate precise relocation. Twenty-four hours post-plating, cells were transfected using Lipofectamine® 2000 (Thermofisher, cat. 11668027) with 1 µg of the LIPA-α-SYN plasmid. Sixteen hours post-transfection, cells were subjected to 12 hrs of blue light stimulation at 0.30 mW/mm^2^. An initial fixation was carried out using a solution of 0.1% glutaraldehyde and 2% paraformaldehyde (PFA) in 0.1 M phosphate buffer (pH 7.4) for 2 hrs at RT. After PBS washes, immunocytochemistry (ICC) was performed as previously described. Fluorescently labeled COS-7 cells expressing LIPA-α-SYN were selected for ultrastructural analysis using a confocal fluorescence microscope (LSM700, Carl Zeiss Microscopy, Germany). The exact coordinates of selected cells were recorded using the dish’s grid. Subsequently, cells underwent a secondary fixation with 2% PFA in 0.1 M phosphate buffer for an additional 2 h. Samples were then washed five times in 0.1 M cacodylate buffer (pH 7.4), post-fixed in 1% osmium tetroxide for 1 h, and rinsed in double-distilled water. Contrast enhancement was performed using 1% uranyl acetate for 1 hr at RT. Dehydration was achieved through a graded ethanol series (2 × 50%, 1 × 70%, 1 × 90%, 1 × 95%, and 2 × 100%), with 3-min incubations at each step. Cells were infiltrated with Durcupan resin (Electron Microscopy Sciences, cat. 14040), progressively mixed with ethanol in ratios of 1:2, 1:1, and 2:1 for 30 min each, followed by two 30-min incubations in 100% resin. Samples were left in fresh Durcupan for 2 hrs before being polymerized overnight at 65°C. Once the resin had fully cured, the glass coverslip was removed by sequential immersion in hot water (60°C) and liquid nitrogen. The previously identified cell was relocated using the recorded grid coordinates. This region was carefully excised from the block, mounted onto a resin stub using acrylic adhesive, and trimmed using a glass knife on an ultramicrotome (Leica Ultracut UCT, Leica Microsystems). Ultrathin sections (50 nm) were cut with a diamond knife (Diatome, Biel, Switzerland) and mounted onto 2 mm single-slot copper grids coated with a Formvar support film. Sections were post-stained with uranyl acetate and lead citrate before imaging with a transmission electron microscope (Tecnai Spirit EM, FEI, The Netherlands) operating at 80 kV and equipped with a digital camera (FEI Eagle, FEI). TEM images and confocal fluorescence images were then manually overlapped to make the final image.

#### In vivo

Brains were sectioned at 50 µm using a vibratome. Sections were immediately mounted in PBS between a microscope slide and a coverslip for rapid fluorescence screening.

Sections containing the highest density of fluorescent neurons in the substantia nigra were selected for tile scanning (63× objective) and optical sectioning (1 µm z-steps) using a confocal microscope. Following confocal acquisition, the ROI was excised in the shape of an asymmetric trapezoid to preserve orientation. Sections were washed three times for 5 min in phosphate buffer (PB, pH 7.4) and postfixed for 1 hr in 2% osmium tetroxide, then post-fixed for 1 hour in 2% osmium tetroxide (Electron Microscopy Sciences, cat. 19152) diluted in 1.5% potassium ferrocyanide (MP Biomedicals, cat. 152560). Samples were rinsed three times in double-distilled water (ddH O), incubated for 20 minutes in 1% thiocarbohydrazide (TCH; MP Biomedicals, cat. 152560), and rinsed again in ddH O. Sections were subsequently incubated for 30 min in 2% osmium tetroxide, following the serial block-face SEM protocol^122^. Dehydration was carried out through a graded ethanol series (30%, 50%, 70%, 90%, 100%), followed by immersion in propylene oxide (Electron Microscopy Sciences, cat. 20401). Samples were then infiltrated overnight in Durcupan resin, sandwiched between ACLAR films, and polymerized at 60°C for 72 hrs. Areas of interest were trimmed into asymmetric quadrangles, mounted onto resin blocks, and sectioned into 50 nm ultrathin slices using an ultramicrotome (EM UC7, Leica Microsystems). Ultrathin sections were collected on single-slot copper grids (Electron Microscopy Sciences, cat. G150-Cu) and imaged using a Tecnai 12 transmission electron microscope (Philips Electronics, Amsterdam, Netherlands) operating at 100 kV. Complete ultrathin sections were manually acquired at the electron microscope. Individual images were stitched using Fiji software to reconstruct the ultrastructural overview of the ROI. Correlation between confocal and TEM datasets was achieved by aligning anatomical landmarks such as blood vessels, myelinated fibers, and the spatial distribution of fluorescently labeled neurons.

### Purification of aggregates and TEM

LIPA-α-SYN aggregates were purified from cells as previously described^20^. Briefly, HEK293T cells stably expressing LIPA-α-SYN^23^ were exposed to blue light for 12 hrs to induce stable aggregation. Cells were collected with a cell scraper after a wash with PBS Ca+ Mg+ in the lysis buffer (PBS 0.05% Triton, supplemented with PMSF, Protease and phosphatase inhibitors). Cells were then lysed with a Dremel tissue homogenizer. LIPA-α-SYN aggregates were clarified, following a first centrifugation at 500 x G for 5 min. Supernatant was then spin at 1000 x G for 10 min. The final supernatant was collected, and protein levels were quantified using BCA (Thermofisher, cat. 23225). LIPA-α-SYN aggregates from cell samples were added to a 200-mesh copper-carbon grid (SPI Supplies, cat. 3520C-FA) for 2 min. The grids were then incubated in 0.1% fish gelatin PBS blocking solution during 10 min, followed by incubation with anti-phospho-synuclein antibody (Fujifilm WAKO, cat. 015-25191) diluted at 1:50 in 0.1% fish gelatin in PBS for 1 h. After rinsing with PBS, the grids were incubated with a secondary antibody goat anti-mouse coupled with 10 nm gold particle (Aurion, cat. 810.022-05) diluted at 1:20 in 0.1% fish gelatin in PBS for 1 h. Then the grids were fixed with 2.5% glutaraldehyde in 1/3 diluted PBS for 5 min, rinsed with distilled water and further stained with 2% acetate uranyl (EMS, cat. 22400-2) for 1 min. The LIPA-α-SYN aggregates were visualized using a transmission electron microscope (FEI Tecnai G2 Spirit Twin 120kV TEM) coupled to a camera (Ultrascan 4000 4k x 4k CCD Camera model 895). Vesicle profiles were classified based on our previous work^20^.

### RNA isolation and quantitative real-time reverse transcription polymerase chain reaction

Total RNA was extracted from HEK293T cells stably expressing either scramble or shRNA constructs using the RNeasy Mini Kit (Qiagen, cat. 74104). RNA was quantified, and for each sample, 1 μg of total RNA was treated with ezDNase™ Enzyme (ThermoFisher, cat. 18091150), and reverse-transcribed using SuperScript™ IV First-Strand Synthesis System from ThermoFisher Scientific (ThermoFisher Scientific, cat. 18091150) according to the manufacturer’s protocol. Quantitative real-time reverse transcription polymerase chain reaction (qRT-PCR) was performed with the QuantiNova SYBR Green PCR Kit using 10 μg/ml of complementary DNA according to the manufacturer’s protocol (Qiagen, cat. 208054) and run on the LightCycler® 96 Instrument (Roche Diagnostics) under the following conditions: PCR initial activation step 2 min at 95 °C, 2-step cycling denaturation 5 s at 95 °C, and combined annealing and extension for 10 s at 60 °C or 50 °C depending on the primer set evaluated, and the qRT-PCR was performed for 40 cycles. Transcript levels of Glyceraldehyde 3-phosphate dehydrogenase (GAPDH) were measured as endogenous controls. Gene expression was analyzed based on the delta CT (ΔCT) approach and normalized to the expression of GAPDH. Primer sequences and melting temperatures are listed in Supplementary Table I.

## Biochemistry

### Western Blot

#### In vitro

Whole cell lysates were collected in 2X Laemmli lysis buffer (0.125 M Tris–HCl (pH 6.8), 20% glycerol, 0.2% 2-mercaptoethanol, 0.004% bromophenol blue, 4% SDS) after being washed twice with dPBS Ca+ Mg+. Samples were then prepared as previously described^20,21^. Briefly, samples were heated at 95 °C for 15 min to allow for DNA denaturation (unless otherwise indicated). Approximately 10 μL of the total protein fraction (corresponding to 35 μg of protein) was loaded per well in a 10% SDS-PAGE gel. The gels were run at 100 V for 90 min prior transfer of the protein to a nitrocellulose membrane (BioRad. cat. 1620112) using wet transfer as previously descrbed^21^. The membranes were then incubated with a blocking buffer (PBS-Tween 0.1%, 3% Fish Gelatin) at RT for 1 hr on a shaker set at a low rotation speed. The membranes were then incubated with primary antibodies (table I) for 1h: anti-mCherry (Abcam, cat. Ab167453), anti-LAMP2 (ThermoFisher, cat. MA1-205), anti-GFP (Abclonal, cat. AE012), anti-α-SYN (BD Laboratories, cat. 610787), anti-beta-actin (ABM, cat. G043), anti-WDR44 (Bethyl, cat. A301-440A), anti-IGF2R (Proteintech, cat. 20253-1-AP), anti-BLTP3A (Proteintech, cat. 24345-1-AP). The membranes were then washed 3x for 10 min with PBS-Tween 0.1% prior to incubation with secondary antibodies matching the host of the primary antibodies using either 680RD or 800CW LiCor antibodies (LiCor, cat. 926–68071). The membranes were then washed with PBS-Tween 0.1% and the signal was acquired before incubating with the next primary antibody. Visualization and quantification were carried out with the LI-COR Odyssey scanner and software (LI-COR Lincoln, NE, USA).

#### Human tissue processing – Saskatchewan cohort

For Western Blotting experiments, proteins were extracted from tissue homogenates, and details on tissue processing are in appended articles and other publications^109^. Briefly, a homogenization lysis buffer containing detergents (0.5% of deoxycholate, 150 mM NaCl, 1% of Triton X-100, 10 mM NaH_2_ PO_4_, 0.5% sodium dodecyl sulphate (SDS) with protease and phosphatase inhibitors and 0.1 mM EDTA), followed by sonication and centrifugation to generate a supernatant corresponding to the detergent-soluble fraction. Using the bicinchoninic acid (BCA) protein assay kit, the protein quantity in the detergent-soluble fractions was determined. Proteins from detergent-soluble fractions were added to Laemmli 5X buffer and heated at 95 °C for 5 min (denatured). The same amount of protein per sample (12 ug) was separated on a 12% SDS-polyacrylamide gel by electrophoresis. The proteins were transferred on polyvinylidene fluoride (PVDF; Cytiva Life Sciences, cat. 45-004-095) 0.45 µm membranes. All membranes were blocked with 5% BSA (BioShop, cat. ALB001) in PBS-Tween 0.1% for 1 hr at RT. For the immunoblots, the antibody against WDR44 (Bethyl Laboratories, cat. A301-440A) was used. As a loading control, the antibody against β-actin (ABM, cat. G043) was used at 1:5000. All incubations were done in Superblock™ blocking buffer (Thermo Fisher Scientific) in PBS containing 0.1% Tween 20 and 0.05% sodium azide. After incubation with a primary antibody, the membranes were washed in PBS-Tween 0.1%, followed by an incubation with a horseradish peroxidase (HRP) anti-mouse (Jackson ImmunoResearch Labs) or anti-rabbit secondary antibody (Jackson ImmunoResearch Labs) at 1:40,000 in PBS containing 0.1% Tween 20 and 1% BSA. The detections were done with Amersham Imager 680 (GE Healthcare Bio-Sciences) following revelation with Luminata (Sigma-Aldrich Millipore), a chemiluminescent HRP substrate. For the analysis of band intensities, the Image Lab software (Bio-Rad) was used.

### Discontinuous Sucrose Density Gradient Fractionation

HEK293T cells, seeded in 60-mm plates, were transiently transfected with the LIPA-α-SYN construct following calcium phosphate transfection. The next day, cells were exposed to blue light at (0.30 mW/mm^2^) for 12 hrs to induce stable aggregation of LIPA-α-SYN. Post light, cells were harvested for membrane fractionation using a sucrose-based membrane preparation buffer. The cells were placed on ice and washed 1-2 times with cold PBS Ca+ Mg+. Cells were then scraped into 450 µL of membrane preparation buffer containing 8% sucrose, 20 mM Tris-HCl (pH 8.0), and 1 mM EDTA. The buffer was freshly prepared by adjusting the pH of Tris-HCl to 8.0 before adding sucrose and EDTA. Cells were then lysed on ice by triturating using a protocol adapted from Sanders *et al.*^123^. The lysates were clarified by centrifugation at 500 × g for 5 min at 4 °C to separate the initial soluble and pellet (debris) fractions. 10% of the lysate was set aside as the total fraction for Western blot analysis. The soluble fraction was then centrifuged at 1000 × g for 10 min at 4 °C. The resulting supernatant and pellet fractions were used for the sucrose Density Gradient. 10% of each of the fractions was retained for Western blot analysis. Before the sucrose gradient, the pellet fraction was resuspended in 450 µL of 0.25 M sucrose prepared in 10 mM Tris-HCl (pH 7.4). For the sucrose gradient fractionation, sucrose solutions of varying concentrations were prepared in 10 mM Tris-HCl (pH 7.4) to form a discontinuous gradient as follows (from bottom to top): 2.0 M sucrose, 500 µL, 1.3 M sucrose, 800 µL, 1.16 M sucrose, 800 µL, 0.8 M sucrose, 800 µL, 0.5 M sucrose, 700 µL, sample 300 µL. Gradients were subjected to ultracentrifugation at 36,000 rpm for 2.5 hrs at 4 °C using a swinging-bucket rotor. Fractions (14 × 280 µL) were collected sequentially from top to bottom. Proteins from each fraction were mixed with 2x Laemmli buffer and analyzed by SDS-PAGE Western blotting. Briefly, soluble fractions were prepared using 5x Laemmli buffer and the insoluble fractions were mixed in equal parts with 2x Laemmli buffer. Protein concentration was quantified by BCA assay, and samples were normalized to 5 µg total protein per well.

### Filter trap assay

HEK293T cells knocked down for WDR44 (WDR44^KD^) or scramble cells were transiently transfected with LIPA-α-SYN plasmid and further used for filter retardation analysis as previously described^20,21^. After transfection, cells were collected at different time points post-illumination with blue light (0.3 mW/mm^2^) as indicated in the figures. Briefly, HEK293T cells were lysed in 250 μL of lysis buffer (PBS-T 0.05% + Protease inhibitor, Phosphatase inhibitor II, Phosphatase inhibitor III and PMSF) per 60-mm plate following two washes with cold PBS Ca+ Mg+. Cells were homogenized using a Dremel tissue homogenizer (BioSpec Products). Cell lysates were sequentially centrifuged at 4 °C (500 x g for 5 min and 1000 x g for 10 min). The supernatant was collected, and 10% of the volume was kept as input sample and diluted in 2x Laemmli, which was checked using Western Blot analysis to confirm equal loading. The clarified supernatant was collected and incubated with SDS to a final concentration of 1.5% for 10 min at RT. Before loading, 0.2 μm cellulose acetate membranes (Sterlitech, cat. CA022005) pre-soaked in PBS containing 1.5% SDS were mounted on top of pre-soaked filter paper on a vacuum manifold. A pre-wash with 100 μL of 1.5% SDS was performed per well, followed by rapid loading of 50 μL of each SDS-treated sample (in triplicate). After sample filtration, wells were washed with 100 μL of 1.5% SDS and the vacuum was applied again. The acetate membrane was then rinsed in Milli-Q water. The immunoblotting was carried out as described in the Western Blot part using an anti-mCherry antibody (table I).

### Immunoprecipitation (IP)

#### Lysosomes Immunoprecipitation (LysoIP)

HEK293T cells stably expressing Lamp1-mGFP-2Strep construct^30^ (gift from Dr. Mahul-Mellier) were generated by infecting cells with lentiviral particles purified from HEK293T cells. Cells were then selected with puromycin at 1 µg/ml for a week. After FACS sorting, cells were seeded in 10-cm plates at 70% confluency and then transfected with either LIPA-α-SYN/LIPA-EMPTY/LIPA-TDP-43, or with either LIPA-α-SYN/ LIPA-α-SYN/LIPA-α-SYN^Δ2-4^/LIPA-α-SYN^Δ2-10^/LIPA-α-SYN^Δ2-18^/LIPA-α-SYN^Δ^^104–140^/LIPA-α-SYN^Δ115–140^/LIPA-α-SYN^Δ121–140^/LIPA-α-SYN^Δ136–140^ following the lipofectamine®3000 transfection protocol. 16 hrs post-transfection, cells were exposed to blue light for a total of 1h. HEK293T cells transfected with the LIPA constructs not exposed to blue light were used as a negative control. One hour post blue light exposure (or not), media was removed, cells were carefully washed once with warm dPBS Ca+ Mg+ to remove any free amines that might react with the crosslinker, and 50 μM disuccinimidyl glutarate (DSG) (Thermofisher, cat. A35392) freshly diluted in warm dPBS Ca+ Mg+ was added to the plates before incubation at 37°C for 30 min under the blue light (light exposure totals to around 1h30) or in the dark. After incubation, dPBS containing DSG was removed, and cells were washed twice in cold dPBS Ca+ Mg+ to remove any remaining DSG. Whole cell lysates were scraped in lysis buffer (PBS-T 0.05% + Protease and phosphatase inhibitor, and PMSF). The lysates were kept on ice for 5 min. Once the cells were swollen, cells were lysed through gentle shearing with a syringe equipped with a 27 G gauge needle (Fisher Scientific, cat. 14-826-87). The progress of cell lysis was monitored by phase contrast microscopy to ensure sufficient lysis, using 25 syringe cycles. Samples were sequentially centrifuged at 4°C (500 x g for 5 min and 1000 x g for 10 min). The supernatant was collected and 10% of the volume was kept as input sample and diluted in NuPage Sample Buffer (4X) containing lithium dodecyl sulfate (LDS) (Thermofisher, Cat. NP0007), supplemented with 2-Mercaptoethanol (5%, final concentration). The remaining supernatant was then used for the LysoIP. Immunoprecipitation was performed using Streptavidin Magnetic Beads (Thermofisher, cat. 88817) following the manufacturer’s instructions and Dong et al^124^. First, a volume of 50 μL (per 200 μL of sample) of Streptavidin beads was transferred to clean microcentrifuge tubes. Tubes were placed on the magnet to separate the beads from the solution. Beads were washed 3 times with Triton lysis buffer. 200 μL of sample supernatant was added to the beads. The beads-sample complex was incubated with rotation for 1 hr at 4°C. The beads were then washed in KPBS buffer (136mM KCl, 10mM KH2PO4 at pH 7.25) 3x by gentle pipetting. After the last wash, the magnetic bead-sample complex was resuspended in 50 μL of KPBS and transferred to a clean tube. To elute the samples, 20 μL of elution Buffer (50 mM glycine at pH 2.8) was used, and 10 μL of premixed NuPAGE LDS Sample Buffer and 2-Mercaptoethanol (5%, final concentration). The samples were then heated for 10 min at 70 °C. Samples were then subjected to immunoblotting as described in the Western Blot protocol.

#### mCherry pulldown

HEK293T cells stably expressing the LIPA constructs (LIPA-α-SYN, LIPA-EMPTY and LIPA-TDP-43), generated as previously described^21,23^, were seeded in 10-cm plates (one per construct) and further used for pulldown experiments. Cells were exposed to blue light for a total of 1.5 h. HEK293T cells stably expressing the LIPA constructs not exposed to blue light were used as a negative control. One hour post blue light exposure or dark condition, cells were treated with DSG and further lysed as described in the LysoIP protocol. Immunoprecipitation was performed using Dynabeads Protein G (Thermofisher, cat. 10004D) following the manufacturer’s instructions. First, a volume of 50 μL (per 200 μL of sample) of dynabeads was transferred to clean microcentrifuge tubes. Tubes were placed on the magnet to separate the beads from the solution. 2 µg of the mCherry antibody (table III) diluted in 200 μL of PBS pH 7.4 with 0.02% Tween-20 was added to the magnetic beads. The beads-antibody complex was incubated with rotation for 30 min at RT. The beads were then washed in PBS pH 7.4 with 0.02% Tween-20 3 x by gentle pipetting. After the last wash, samples were added to the magnetic bead-antibody complex, the samples were incubated with rotation for 16 hrs at 4 °C to permit antigen binding to the magnetic bead-antibody complex. After the incubation time, the sample tubes were placed on the magnet, and the supernatant was removed. The magnetic bead-antibody-sample complex was washed 3 x by gentle pipetting using 200 μL of PBS pH 7.4. The magnetic bead-antibody-sample complex was resuspended in 100 μL of PBS pH 7.4 and transferred to a clean tube. To elute the samples, 20 μL of elution Buffer (50 mM glycine at pH 2.8) was used, and 10 μL of premixed NuPAGE LDS Sample Buffer and 2-mercaptoethanol (5%, final concentration). The samples were then heated for 10 min at 70 °C. Samples were then subjected to immunoblotting as described in the Western Blot protocol.

## Acknowledgements

We thank the Bio-EM Facility at for access to electron microscopes and manual CLEM reconstructions. This work was supported by the Canadian Institutes of Health Research (CIHR; FRN 162400) to A.O., the Natural Sciences and Engineering Research Council (NSERC; RGPIN-2023-05581) to A.O and Society Parkinson Canada to A.O. A.O. was supported by Junior 2 salary awards from the Fonds de Recherche du Québec – Santé (FRQS) and la Société Parkinson du Québec. C.J.W., and C.I. are supported by the Intramural Research Program of the National Cancer Institute (contract #HHSN26120080001E). E.B., and M.-H.C. received funding from the European Research Council (ERC) under the European Union’s Horizon 2020 research and innovation program (Grant agreement No. #951294 to EB) and financial support from the French government in the framework of the University of Bordeaux’s IdEx “Investments for the Future” program/GPR BRAIN_2030. F.Ci. receives funding, in part, from the Canadian Institutes of Health Research E.L., C.T. and M.T. are supported by a doctoral scholarship from the Fonds de recherche du Québec-Santé (FRQS). N.R. and U.D. are supported by the Neuroscience Support grants from the National Institutes of Health (NIH) (AG085401 to N.R., NS099328 to U.D. and NS133979 to N.R. and U.D.). A.L.M. was supported by EPFL and the Fundation Bru. F.Ca. and A.J.R. were supported by Parkinson Canada (2017-1110). E.A.F. is supported by a grant from the Canadian Institutes of Health Research (PJT-195804) and by a Canada Research Chair (Tier 1) in Parkinson’s disease.

## Ethics declaration

H.A.L. is the co-founder and chief scientific officer of ND BioSciences, Epalinges, Switzerland, a company that develops diagnostics and treatments for neurodegenerative diseases. The other authors declare no competing interests.

## Contributions

Conceptualization, M.T. and A.O.; Data curation, M.T. and R.S.; Formal analysis, M.T., R.S., C.I., E.L., and F.Ca.; Funding acquisition, A.O.; Investigation, M.T., R.S., M.B., C.I., A-L.M., C.V.L.D., E.L., W.I., A.Ri., E.D.C-P., C.T., M-H.-C., and F.Ca.; Methodology, M.T., R.S., and; Project administration, M.T., R.S. and A.O.; Resources, S.G., E.B., A.Ra., F.Ci., M.P., E.A.F., N.R., U.D., C.W., H.A.L. and A.O. Supervision, M.T., R.S., and A.O.; Validation, M.T., R.S., M.B., E.L., W.I., F.Ca.; Visualization, M.T.; Writing – original draft, M.T. and A.O.;Writing – review and editing, M.T., R.S., A.L.M., S.G., E.B., F.Ci., M.P., E.A.F., N.R., U.D., F.Ca., C.W., H.A.L., and A.O.

---

**Extended Data Fig. 1:**
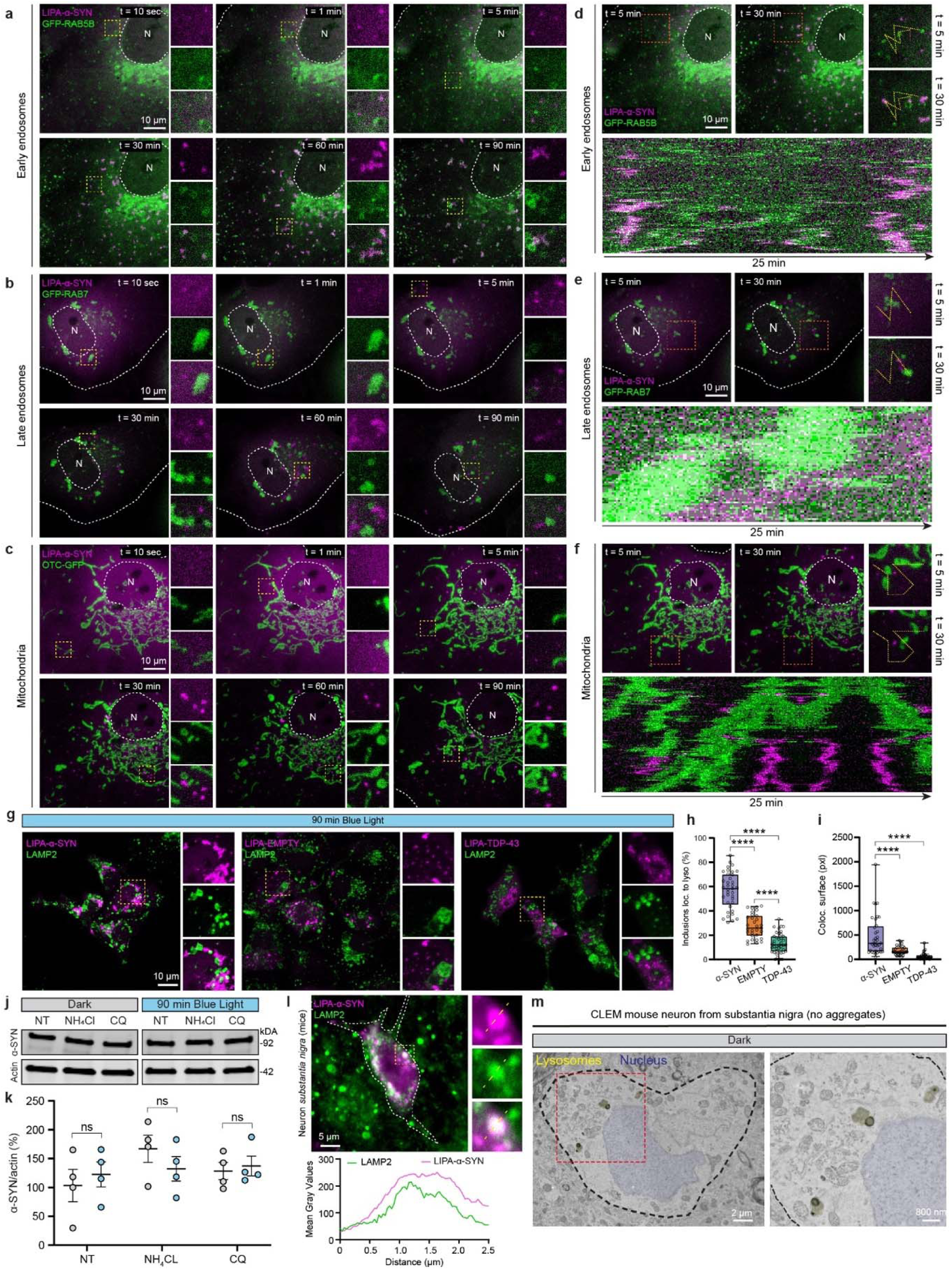
Interaction with lysosomal membrane is specific to nascent LIPA-**α**-SYN aggregates and is independent of the autophagy-lysosomal degradation pathway. **a**, **b**, **c**, Representative live-cell confocal microscopy images of COS-7 cells expressing LIPA-α-SYN and either GFP-RAB5B (early endosomes (**a**)), GFP-RAB7 (late endosomes (**b**)), and OTC-GFP (mitochondria (**c**)), showing the absence of α-SYN accumulation at the organelles membranes after blue light exposure (after 10 sec, 1, 5, 30, 60 and 90 min). The small panels on the right show magnified views of the region highlighted by the yellow box. Scale bars, 10 µm. N = nucleus. **d**, **e**, **f,** Kymographs demonstrating absence of co-traffic or interactions between LIPA-α-SYN aggregates and early endosomes in **d**, late endosomes in **e**, and mitochondria in **f**. Scale bars, 10 µm. N = nucleus. **g**, Representative confocal images of COS-7 cells transfected with either LIPA-α-SYN, LIPA-EMPTY or LIPA-TDP-43 (magenta) and further stained for endogenous LAMP2 (green), showing only the accumulation of LIPA-α-SYN aggregates at the lysosomal membrane. The small panels on the right show magnified views of the region highlighted by the yellow box. Scale bars, 10 µm. **h**, **i**, Quantitative analysis of LIPA aggregate-lysosome association (**h**), as well as the colocalization surface between lysosomes and the aggregates in 3D (**i**). Both quantifications are displayed as box and whiskers (n=3, ∼ 40 individual cells quantified, one-way ANOVA Tukey’s multiple comparison tests). **j**, Representative Western blots showing no change in LIPA-α-SYN levels in HEK293T cells after autophagy inhibition using NH Cl or chloroquine (CQ)). **k**, Quantification of α-SYN band intensity after normalization to actin, displayed as mean ± SEM (n=4, 2-way ANOVA with Dunnett’s multiple comparisons tests). **l**, Representative confocal image of a mouse dopaminergic neuron overexpressing AAV-LIPA-α-SYN and exposed to blue light (1 h/day) for 7 days and further stained for endogenous lysosomes (LAMP2, green). The small panels on the right show magnified views of the region highlighted by the yellow box. The intensity profile of the yellow dotted line is displayed below to show the overlap between the two signals. Scale bar, 5 µm. **m**, Representative TEM micrographs (CLEM protocol) of a mouse dopaminergic neuron overexpressing AAV-LIPA-α-SYN but not exposed to blue light. Neuron displays a normal lysosomal phenotype. The small red panel was used for the magnified view represented on the right. Scale bar, 2 µm (left) and 800 nm (right). Each experiment was independently replicated at least three times, and results were reproducible across replicates. *\* p* < 0.05, *\*\* p* < 0.01, *\*\*\* p* < 0.001, *\*\*\*\* p* < 0.0001.

**Extended Data Figure 2:**
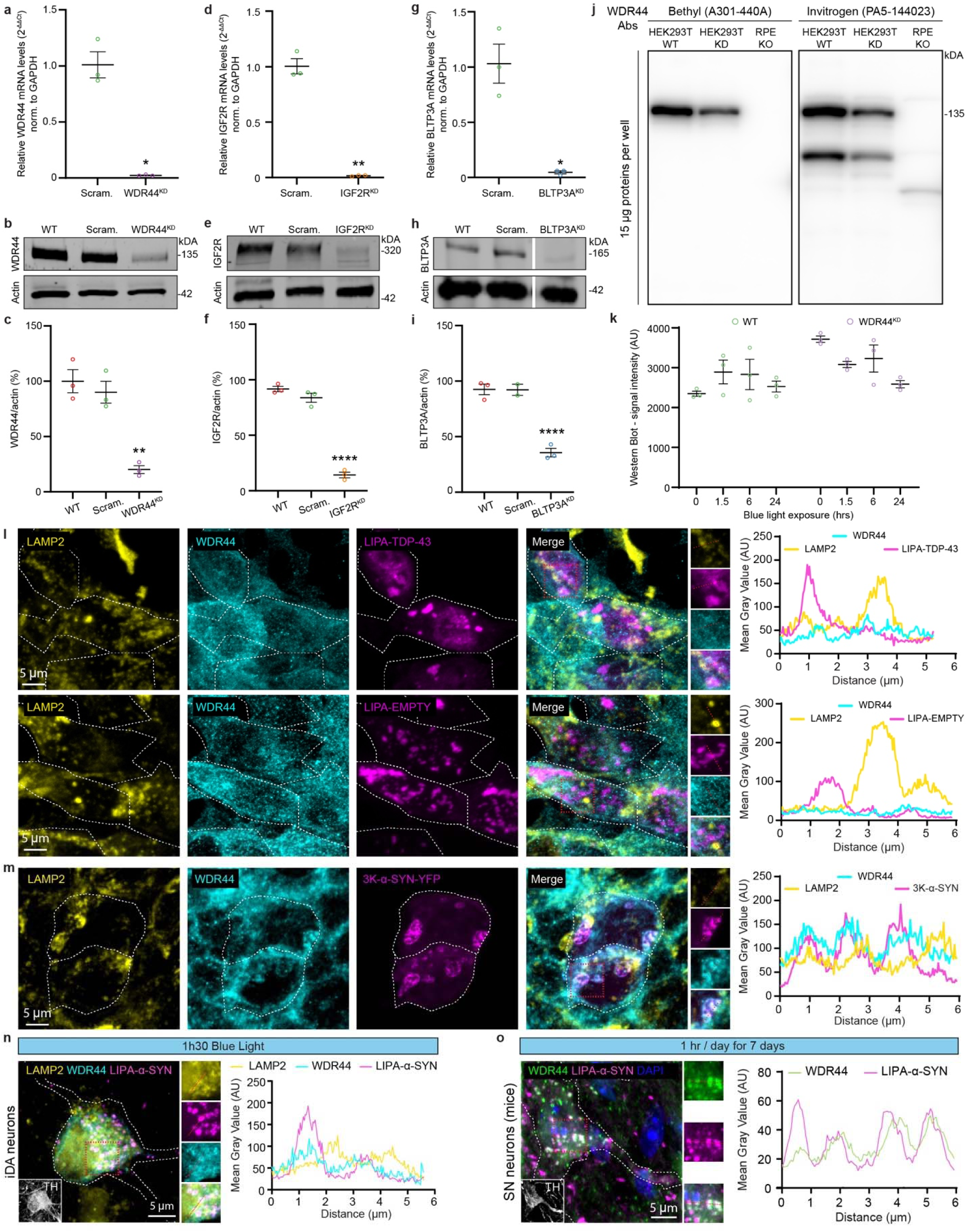
Validation of WDR44 knockdown and confirmation of WDR44 colocalization with **α**-SYN *in vivo* and neuronal culture. **a**, **b**, **c**, Knockdown validation for WDR44^KD^ induced by shRNA at the mRNA level (**a**) and at the protein level with a representative Western blot (b) and its relative quantification (c, scatter dot plot represented as mean ± SEM, n=3). **d**, **e**, **f** Knockdown validation for IGF2R^KD^ induced by shRNA at the mRNA level (d) and at the protein level with a representative Western blot (e) and its relative quantification (f, scatter dot plot represented as mean ± SEM, N=3). **g**, **h**, **i** Knockdown validation for BLTP3A^KD^ induced by shRNA at the mRNA level (g) and at the protein level with a representative Western blot (h) and its relative quantification (i, scatter dot plot represented as mean ± SEM, N=3, 2-way ANOVA with Dunnett’s multiple comparisons test comparing all conditions to the WT (WB)). **j**, Representative Western blot membrane showing the specificity of the antibodies targeting WDR44 used in this study (Bethyl for biochemistry, Invitrogen for IF). Samples loaded were HEK293T lysates (WT or shRNA KD) or RPE-E1 knockout for WDR44 (through CRISPR-KO^53^). **k**, Quantification of total mCherry protein levels for the filter retardation assay related to Fig. 3c and showing equal loaded protein levels (scatter dot plot represented as mean ± SEM, n=3, 2-way ANOVA with Dunnett’s multiple comparisons test). **l**, Representative confocal images showing HEK293T transfected with LIPA-TDP-43 (top panel) or LIPA-EMPTY (bottom panel), exposed to 90 min blue light and stained for endogenous LAMP2 (yellow) and WDR44 (cyan). The small panels on the right show magnified views of the region highlighted by the red box. Intensity profiles (right) are displayed over the red-dotted line in the small panels. Scale bar, 5 µm. **m**, Representative confocal images showing HEK293T transfected with 3K-α-SYN exposed to 24 hrs of doxycycline and stained for endogenous LAMP2 (yellow) and WDR44 (cyan). The small panels on the right show magnified views of the region highlighted by the red box. Intensity profiles (right) are displayed over the red-dotted line in the small panels. Scale bar, 5 µm. **n**, Representative confocal images showing iDA neurons transfected with either LIPA-α-SYN exposed to 90 min blue light and stained for endogenous TH (white), LAMP2 (yellow) and WDR44 (cyan). The small panels on the right show magnified views of the region highlighted by the red box. Intensity profiles (right) are displayed over the red-dotted line in the small panels. Scale bar, 5 µm. **o**, Representative confocal images showing mouse dopaminergic neurons infected with an AAV-LIPA-α-SYN, exposed to the blue light for 1h/day for 7 days and stained for endogenous TH (white) and WDR44 (green). The small panels on the right show magnified views of the region highlighted by the red box. Intensity profiles (right) are displayed over the red-dotted line in the small panels. Scale bar, 5 µm. Each experiment was independently replicated at least three times, and results were reproducible across replicates. * p < 0.05, ** p < 0.01, *** p < 0.001, **** p < 0.0001.

**Extended Data Figure 3:**
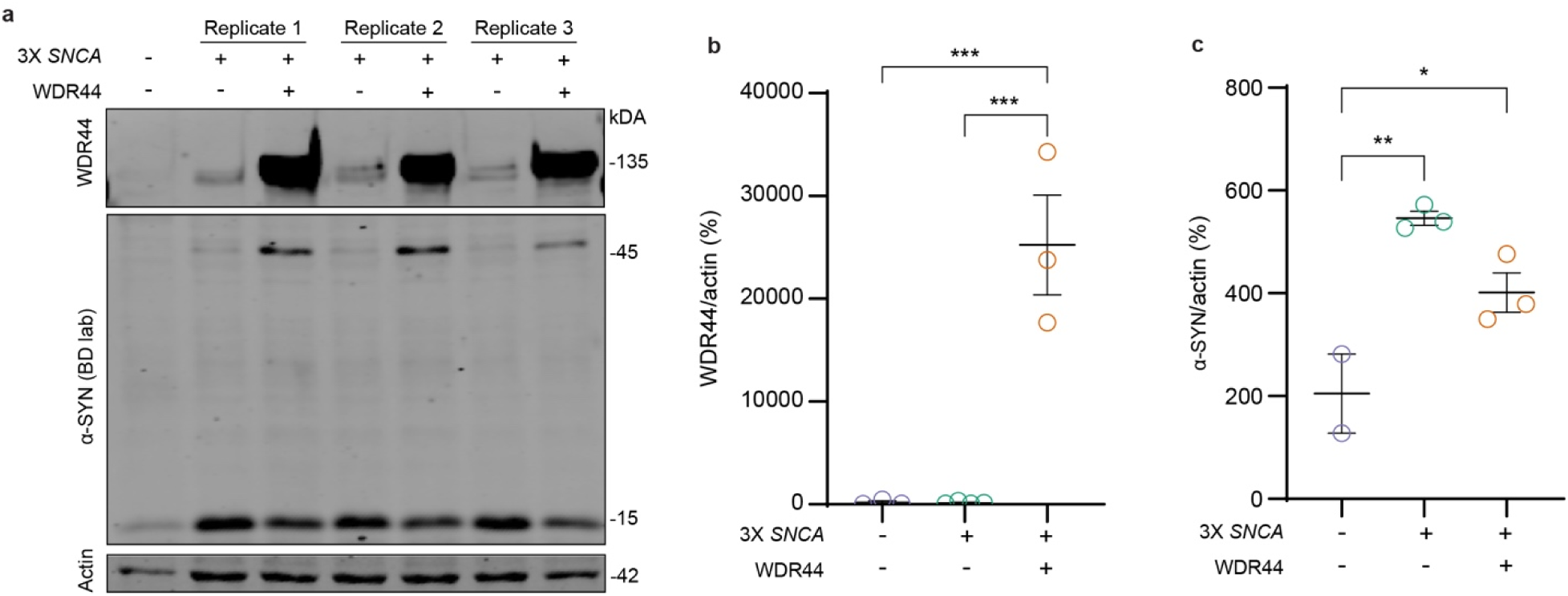
Validation of the 3X *SNCA* hiPSC cell line **a**, Representative Western blot confirming piggyBac-driven WDR44 overexpression in hiPSC-Ngn2-derived neurons (WT or 3X *SNCA* genetic background) after 10 days in culture (with endogenous WDR44 for reference). **b**, Quantification of WDR44 protein levels normalized to actin (scatter dot plot, mean ± SEM; n=3; One-way ANOVA with Tukey’s multiple comparisons test). **c**, Quantification of α-SYN protein levels normalized to actin, confirming the 3X *SNCA* phenotype (scatter dot plot, mean ± SEM; n=3; One-way ANOVA with Tukey’s multiple comparisons test). Each experiment was independently replicated at least three times, and results were reproducible across replicates. *\* p* < 0.05, *\*\* p* < 0.01, *\*\*\* p* < 0.001, *\*\*\*\* p* < 0.0001.

