## Supplementary figures and images for "WDR44 drives de novo α-synuclein aggregation at the lysosomal membrane and promotes neuronal dysfunction in Parkinson’s Disease"

### Suppl Figure 1

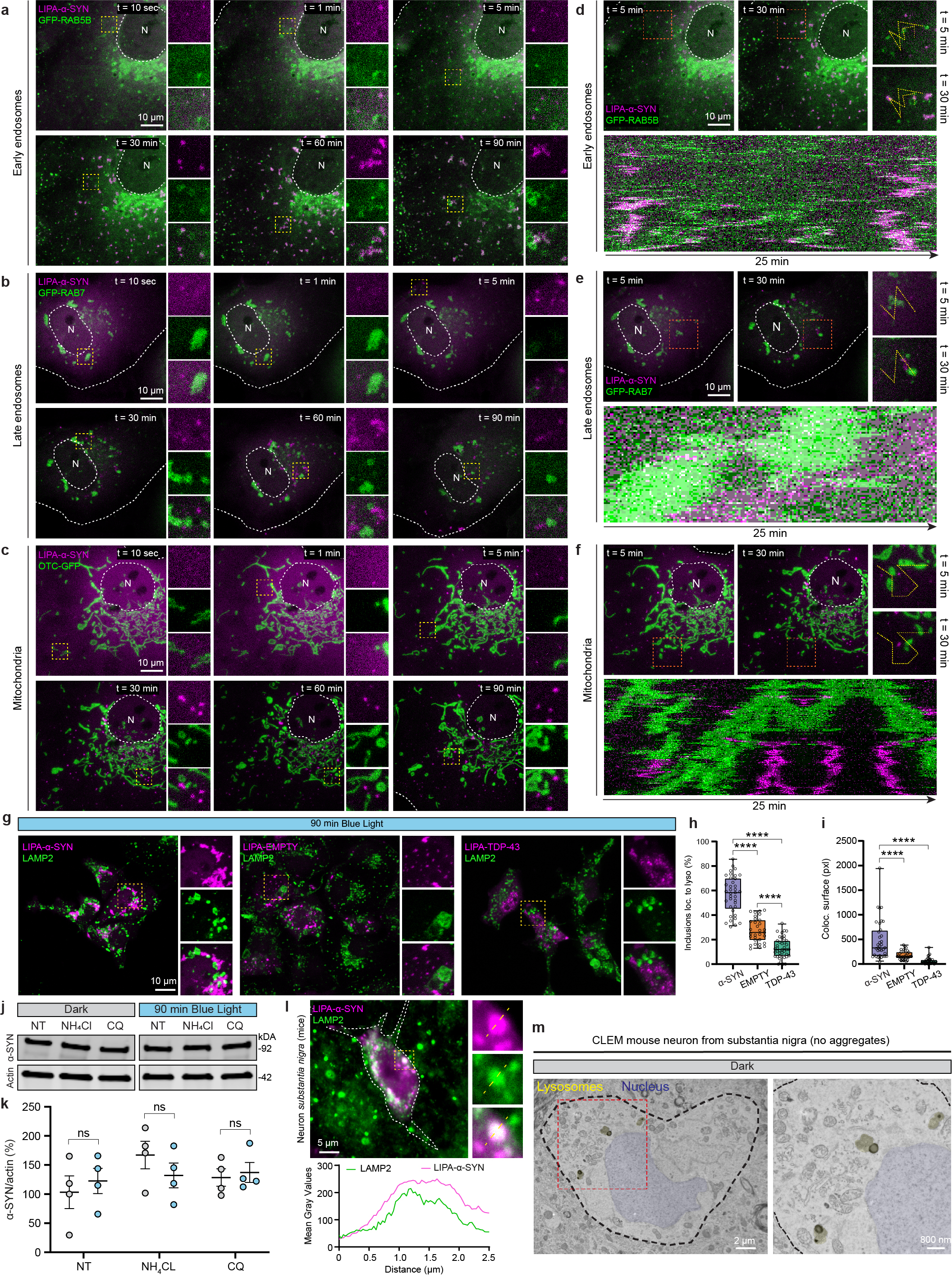

### Suppl Figure 2

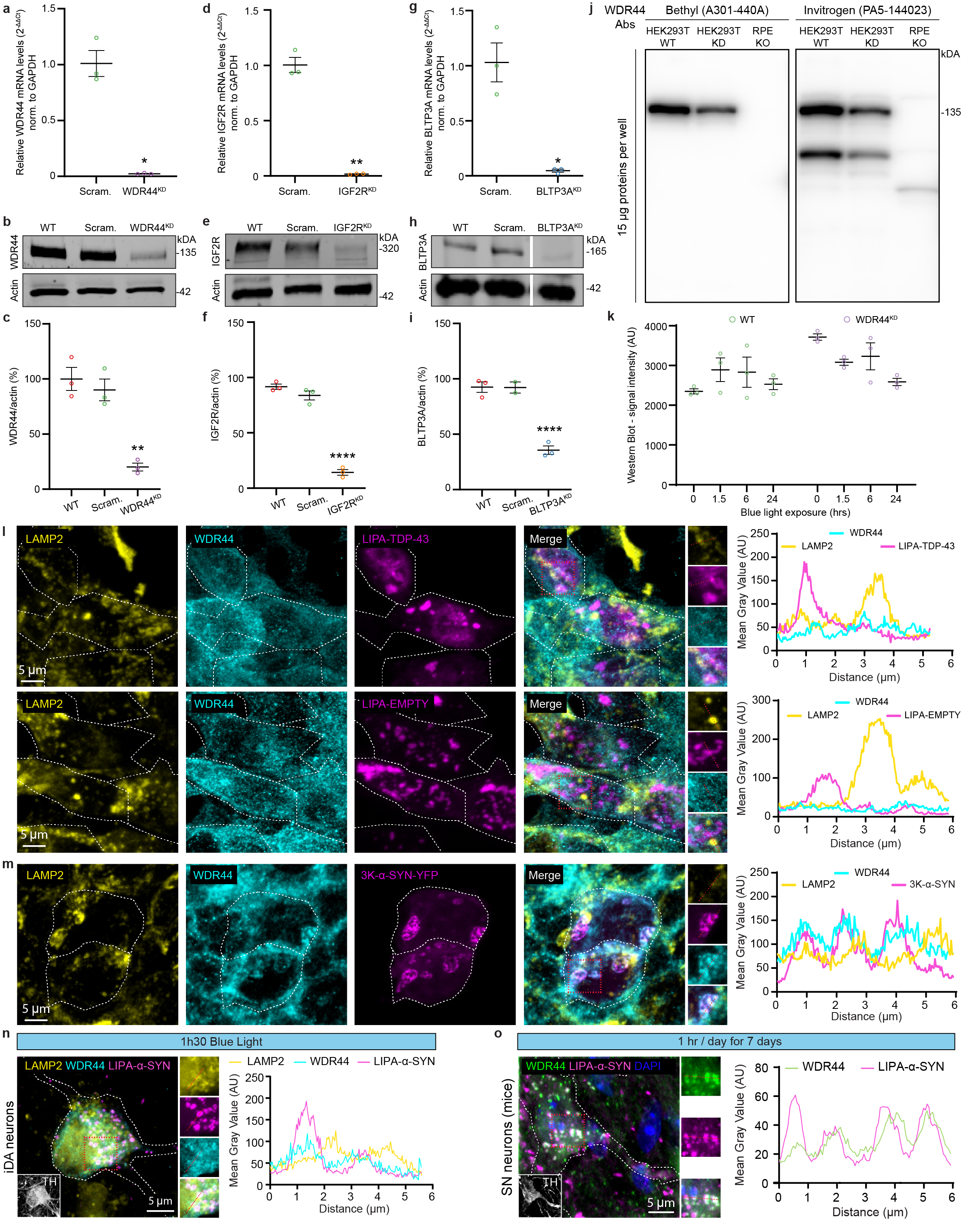

### Suppl Figure 3

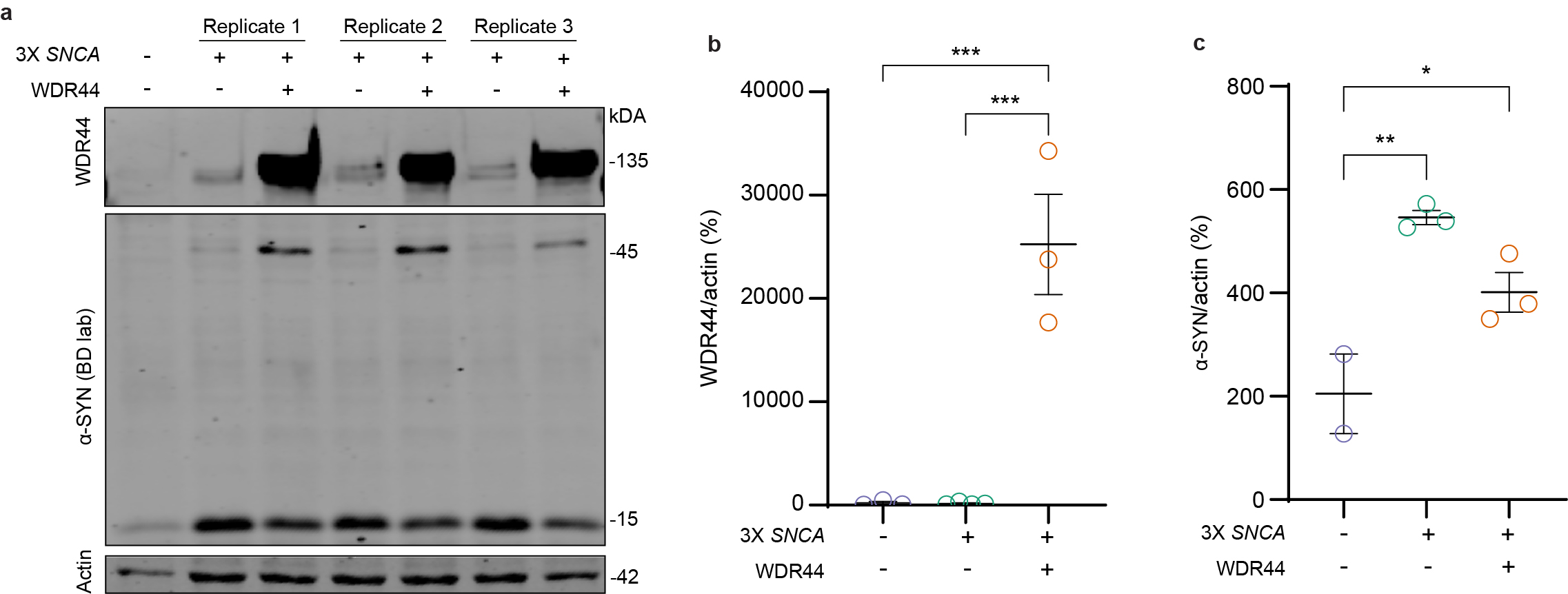
